# Discovering compositional, interpretable structure in social behavior

**DOI:** 10.64898/2026.11.03.764108

**Authors:** Yue Wan, Sai Koukuntla, Ahmed El Hady, Mikio Aoi, Adam S. Charles

## Abstract

Quantitative assessment of behavior is vital for rigorous analysis of how animals interact in the world and how behavior is generated from brain activity. Recent advances in automated pose detection have made possible data-driven discovery of movement action sequences. While many algorithms have been developed for this purpose, their categorical nature, i.e., the assumption that behavior is one action at a time, prohibits expressive quantification of social behavior. Such behavior is combinatorial and compositional, with multiple animals performing actions that may be either independent or coordinated over time. Here we show how developments in decomposed linear dynamical systems (dLDS) can instead represent action-generating dynamics as a signed, graded mixture of shared linear operators acting directly on keypoint positions. We demonstrate our approach across both single-mouse and two-mouse datasets, comparing against a state-of-the-art switching model, keypoint-MoSeq. In single-mouse data, we find that a composition of three operators can capture the full repertoire of 21 keypoint-MoSeq syllables, showing that even for a single animal, the action space is more parsimoniously described by a continuous subspace than clustered motifs. In a two-mouse resident–intruder assay, we identify, without behavioral labels, separate primitives for self-action, inter-animal-geometry, and social-action. The latter couples an animal’s action to its partner’s relative bearing, enabling the action to incorporate the social geometry in which it occurs. dLDS further decodes human-labeled social behaviors (i.e., “attack”, “investigation”, “mount”, and “other”) more accurately than behavior-selective keypoint-MoSeq syllable sets using fewer units. Together, these results demonstrate that compositional, time-varying linear dynamics can provide a path towards flexible and interpretable modeling of action-level behavior for both individual and social behavior.

## 1 Introduction

Quantitative measures of behavior are central to understanding how animals interact with the world and how neural activity relates to behavior. Such measures depend on a choice of behavioral units and on how those units combine. Movement recurs with enough regularity to support a description in terms of units [Tinbergen, 1951, Dawkins, 1976, Baerends, 1976], whose combination proceeds along two axes: temporal and configural. Along the temporal axis, sub-movements combine into sequences and then into longer routines [Lashley, 1951, Simon, 1962, Dawkins, 1976]. Along the configural axis, behavior at each instant comprises several co-occurring components: a turn blends orientation, head movement, and forward translation in proportions that vary from moment to moment [Bernstein, 1967]. In multi-animal contexts, an animal’s action also depends on its geometric relationship to other animals, making relative orientation and distance potentially as important as the action itself [Tinbergen, 1953, Hinde, 1976]. An accurate quantitative account of behavior requires both the temporal and configural axes.

Markerless pose tracking provides the necessary pose information needed to develop such a model, producing dense, continuous trajectories of body configuration across species and for multiple animals [Mathis et al., 2018, Pereira et al., 2022, Biderman et al., 2024]. The remaining challenge is to recover from these keypoint trajectories the higher-order structure that constitutes actions and behaviors. Accomplishing this task requires defining quantitative models that identify recurring temporal and configural patterns. A key design choice for these models is whether to describe behavior at the action level as categorical, i.e., that any time epoch can be considered as one (and only one) of several actions, or as continuous and compositional, i.e., that there are features of actions that define an “action manifold”.

The former paradigm has received the most attention in recent years, with multiple methods developed to assign a series of poses to a set of actions, including switching dynamical systems models [Wiltschko et al., 2015, Weinreb et al., 2024] and deep learning approaches [Hsu and Yttri, 2021, Luxem et al., 2022, Berman et al., 2014], in both supervised [Tillmann et al., 2024] and unsupervised [Wiltschko et al., 2015, Weinreb et al., 2024, Hsu and Yttri, 2021] settings. These models assign each frame to one categorical mode, segmenting a recording into temporal sequences of these modes. The modes themselves can be either class labels, e.g., for supervised deep learning [Tillmann et al., 2024], or autoregressive linear dynamical operators defined over an animal’s egocentric frame, e.g., for unsupervised switching linear dynamical systems [Weinreb et al., 2024].

Along the configural axis, these categorical models of actions lack two kinds of freedom. The first arises within a single animal: a “syllable” can describe coordinated movement across body parts, but only as one invariant pattern. Only a single syllable can be active at a time, indicating a categorical action, and the model cannot freely interpolate between actions or compose complementary actions. For example, such categorical models require three distinct syllables to describe a left turn, grooming, and a left-turn-while grooming. Any combination of co-occurring behaviors then need to be assigned their own, new syllable rather than compose already available actions in new ways.

A second, related, limitation pertains to modeling social behavior between multiple individuals. When more than one animal is being observed, there can be times when the animals interact and times when the animals act independently. Trying to classify the full configuration of both animals with one categorical label per frame becomes combinatorial in nature. In practice, applying categorical models to dyadic recordings can produce various errors. Dynamical systems models, such as keypoint-MoSeq, can track the presence of a conspecific but do not map onto specific social behaviors, [Ritter et al., 2025, Weinreb et al., 2026]. Deep learning methods, such as A-SoiD [Tillmann et al., 2024] can capture frames when both animals are engaged in a shared action but cannot explicitly disentangle the concurrent actions of individual animals.

Here we take an alternative approach where we instead treat actions as features that can combine to define a continuous space of behaviors and more naturally extend to multiple animals. To achieve this, we introduce two techniques that provide the degrees of freedom missing from other approaches. First, we adopt decomposed Linear Dynamical Systems (dLDS), in which the effective linear operator at each instant is a sparse, signed combination of a few stereotyped operators [Mudrik et al., 2024]. Our model expresses actions at each time point as a graded combination of simultaneously active components, creating a continuous rather than discrete parameterization of the configuration space. Because the operators act directly in kinematic feature space, their effects can be deduced from directly analyzing the corresponding values in the dynamical system matrix entries. Second, in social settings, we supplement the pose space with the geometry between the two animals, specifically relative distance and orientation [Berezhnoi et al., 2025, Segalin et al., 2021], so that operators can capture any combination of an animal’s own action and its relationship to a partner. Using this expanded coordinate system, the dLDS decomposition can more readily capture multiple animals since each action can be restricted to one animal’s configuration while being simultaneously active, but also capture social actions by combining both animals’ configuration in the same action atom.

We present our composite behavioral dynamics model (Section 2.1) before applying it to single-mouse and dyadic recordings. In a single animal recording, we demonstrate how our approach identifies a continuous behavioral space that categorical models artificially separate into mutually exclusive states. In social dyads, without behavioral labels, we show that our dLDS model naturally discovers separate operator families for self-action, partner-geometry, and social-action, the latter of which resolves relative distance and orientation. This unsupervised separation of different interaction types allows us to analyze and interpret each operator in the context of distinct behaviors. We show that the operator coefficients decode human-labeled social behavioral labels (“attack”, “investigation”, “mount”, and “other”) more accurately and with fewer units than behavior-selective syllable sets learned by state-of-the-art categorical models.

## 2 Results

### 2.1 Decomposing behavior into a composition of actions

Our model, like other automated behavior identification models, infers an animal’s behavior from pose tracking keypoints. Because the identity of an action is often invariant to absolute position and heading, we first center and rotate the tracked keypoints in absolute coordinates, denoted ***y***_*t*_, into the animal’s own heading frame (Fig. 1a,b). This process leaves an egocentric feature vector of keypoint positions, which we denote as ***x***_*t*_. Because these coordinates are low-dimensional and anatomically labeled, the dynamics can be modeled on them directly to produce interpretable operators whose entries describe couplings between named body parts.

**Figure 1:**
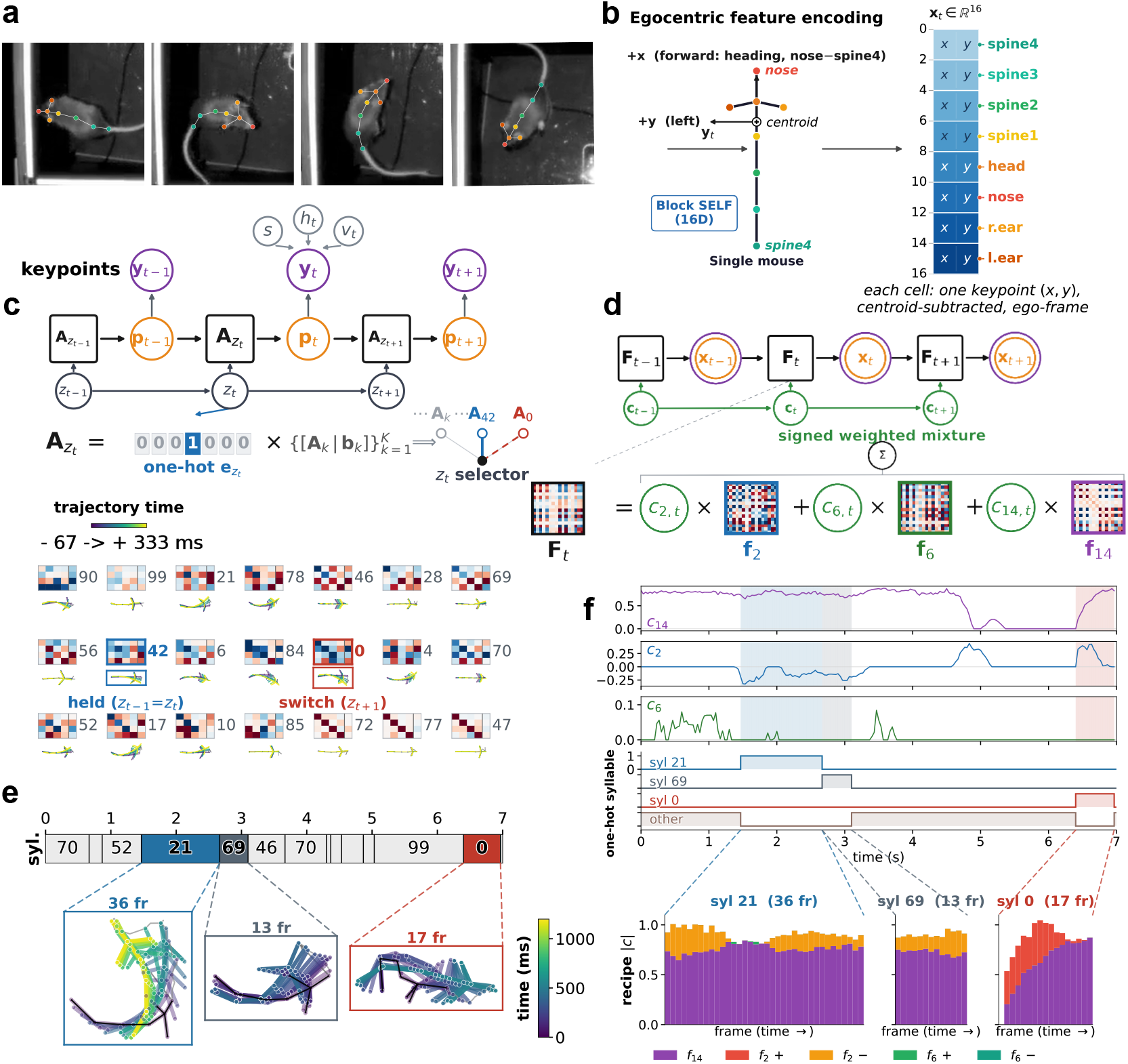
Keypoint-MoSeq switches among syllables where dLDS reweights three operators. **a)** Example frames from a mouse moving in an open field, with eight tracked keypoints overlaid. **b)** Perframe egocentric encoding. Coordinates for the eight keypoints are defined relative to their centroid such that the spine4–nose heading axis points along +*x* and the animal’s left side toward +*y*. **c**,**d)** keypoint-MoSeq and dLDS models fitted to the same five sessions. Box, the dynamics unit. Orange node, the latent state it acts on. **c)** Top, *k*-step dynamics. At each frame, the dynamics follow a single syllable selected from a categorical set (Eq. (2)). Bottom, dynamics matrices and onset-aligned pose trajectories (median over all onsets, −67 to +333,ms, aligned on the onset frame’s position and heading) of the syllables used in more than 5% of frames, out of 72 learned syllables. **d)** Top, one-step dynamics. Each frame’s dynamics are a linear mixture of learned operators (Eq. (3)). Bottom, operator matrices selected as in c, out of 15 learned operators, with usage measured as the share of the pooled |*c*_*m,t*_| mass. **e)** Keypoint-MoSeq assigns one mutually exclusive syllable to each frame of the 7-s trajectory, highlighting syllable 21,69 and 0 bouts. **f)** Signed dLDS coefficient traces over the same interval in e), with the one-hot keypoint-MoSeq sequence below.

Dynamical systems approaches often model the temporal evolution of the state as a time-varying linear system

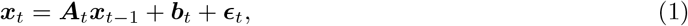

where ***A***_*t*_ is an *d* × *d* matrix that describes how the current state evolves between timepoints, ***b***_*t*_ is a possible offset/input vector, and ***ϵ***_*t*_ is the model error vector. The model parameters (***A***_*t*_ and ***b***_*t*_) are time-dependent because the keypoint dynamics change over time based on what actions the animal is performing. Thus additional constraints or assumptions need to be made to make the model inference possible. In particular, learning a different ***A*** and ***b*** at each point requires fitting *N* ^2^ + *N* variables with only *N* equations. This model is thus extremely over-parameterized without additional constraints.

A recent, and currently state-of-the-art, categorical method that takes the dynamical systems approach— keypoint-MoSeq—makes three modeling choices: 1) the dynamics are modeled in a latent space derived from keypoint positions, instead of in the keypoints themselves 2) each ***A***_*t*_ is drawn from a finite set of *N* possible ***A***’s; and 3) the map at each frame acts on its previous *L* = 3 states.

In keypoint-MoSeq the pose state is a latent variable vector, denoted ***p***_*t*_, which is inferred from the keypoint positions. keypoint-MoSeq reconstructs ***y***_*t*_ from ***p***_*t*_ through a basis ***B***, which is initialized by principal components and held fixed during fitting [Weinreb et al., 2024]. The syllables learned by keypoint-MoSeq therefore operate on points in this *d*-dimensional latent space. ***p***_*t*_ is estimated jointly with the syllables, the animal’s centroid (***v***_*t*_) and heading (*h*_*t*_), and a per-frame, per-keypoint noise scale (*s*_*t,k*_) that absorbs tracking error (Fig. 1c). The joint estimation makes egocentric alignment a central part of the model. The dynamics over ***p***_*t*_ are modeled as an autoregressive process whose parameters change over time (Fig. 1c) as

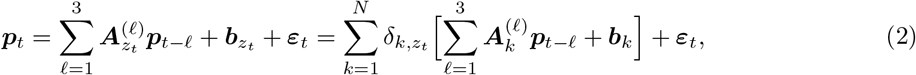

where 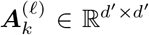 is the dynamics matrix at lag *l* and 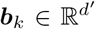 the bias vector, the discrete index *z*_*t*_ selects one dictionary entry 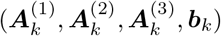 out of *N* syllables, and 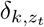 is the Kronecker delta, which allows us to rewrite the selection as a weighted sum over the whole dictionary, with the weight vector confined to the one-hot vertices {***e***_1_, …, ***e***_*N*_}. A sticky prior *κ* then makes switches infrequent, since animals presumably do not oscillate rapidly between actions on single-frame timescales (Supplementary Table S1).

In contrast to keypoint-MoSeq, dLDS fits the dynamics directly on ***x***_*t*_, normalized so that keypoints with larger values (e.g., tail vs. ears) do not dominate the fit. It therefore forgoes both the fixed pose basis ***B*** and the per-keypoint noise scales *s*_*t,k*_ that keypoint-MoSeq uses to separate tracking error from pose dynamics.

Our model also describes the temporal evolution of the state using a linear operator at each timepoint. However, rather than choosing only one of a fixed set of operators per timepoint, we define a linear decomposition of the dynamical system from *M* shared dictionary elements ***f***_*m*_ ∈ ℝ^*d×d*^ (Fig. 1d):

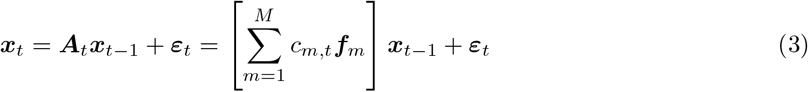

where *d* is the dimensionality of the egocentric feature, *c*_*m,t*_ denotes how strongly dictionary element ***f***_*m*_ impacts the dynamics at time *t*, and ***ε***_*t*_ is again the model error term. There is no bias term. Compared to the keypoint-MoSeq model in Eq. (2), our model describes the dynamics of the egocentric keypoint coordinates directly, rather than the dynamics of an inferred latent state at multiple lags. Our model also uses continuous sparse coefficients *c*_*m,t*_ ∈ ℝ rather than the Kronecker delta 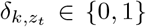 from keypoint-MoSeq. This lifts the constraint that only one action can occur at each time point. To keep this more expressive model interpretable, we assume that only a few actions co-occur relative to the full action repertoire, and the sparsity of *c*_*m,t*_ encodes this assumption.

To differentiate categorical dynamics from composite dynamics, we will adopt the terminology where the keypoint-MoSeq entries are called *syllables* and the dLDS dictionary elements ***f***_*m*_ are called *operators*. Moreover, we will reserve the term *action* for the behavioral level both dictionaries are meant to describe.

We note that ***c***_*t*_, while sparse, can activate several modes at once. For example, dLDS could represent “left-turn-while-grooming” as an appropriate time-varying coactivation of the “left-turn” and “grooming” operators, while keypoint-MoSeq would instead have to learn three syllables for each of “left-turn,” “grooming,” and “left-turn-while-grooming.” If ***c***_*t*_ = [*c*_1,*t*_, …, *c*_*M,t*_] were constrained to permit only a single non-zero element at a time, the dLDS model would reduce to a switching model that defines a single configurational pattern over time, similar to keypoint-MoSeq. In the case where more than one element of ***c***_*t*_ is non-zero, the displacement of each keypoint between two frames would be the weighted sum of the displacements each active operator would produce on its own. The operators thus represent the configurational axis decomposition (entries of each ***f***_*m*_ describe how that operator affects keypoint positions); and *c*_*m,t*_ represents the temporal decomposition by describing how long (*t*), at what strength (|*c*_*m,t*_|) and in which direction (sign(*c*_*m,t*_)) the given configuration’s dynamics persist. The ability to represent the dynamics at a single time point with more than one operator allows our dLDS approach to more readily disentangle components of behavior along both the temporal and configurational axes. Separately, because the decomposition is linear, each ***f***_*m*_ applies the same linear map wherever it appears, so its contribution depends only on the current pose.

The same principles extend to dyads. Dyads’ ***x***_*t*_ concatenates each animal’s egocentric pose with the distance between them and relative orientation, so the learned operators can describe interactions between an animal’s keypoints and its geometric relation to its partner. Because the coordinates are grouped, an operator’s block structure reports which of these it acts on.

### Model learning and constraints for interpretability, identifiability, and efficiency

We fit 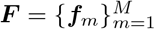 and 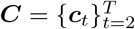 by minimizing

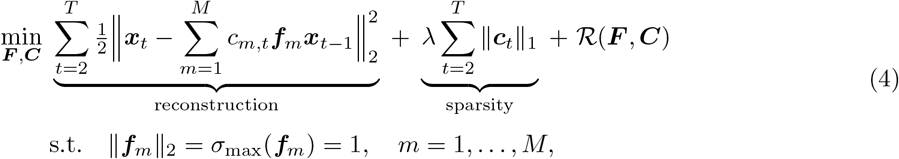

where *σ*_max_(***f***_*m*_) is the largest singular value of ***f***_*m*_, *c*_*m,t*_ is unconstrained in sign, and ℛ includes two further penalties—temporal smoothness to prevent rapid artifactual switching in *c*_*m,t*_’s and pairwise operator decorrelation to avoid redundancy in the dictionary (Methods, Eq. (13)).

The L1 penalty encourages identifiability: we set *M* larger than the state dimension *d*, so the reconstruction accuracy term alone is underdetermined and admits an infinite set of equally good solutions, many of which are not individually interpretable. The L1 penalty favors minimal-support solutions within that set.

The unit-norm constraint removes a scaling degeneracy between *f*_*m*_ and *c*_*m,t*_. The reconstruction depends on the product of ***f***_*m*_ and *c*_*m,t*_, so rescaling (***f***_*m*_, *c*_*m,t*_) ↦ (*α****f***_*m*_, *c*_*m,t*_*/α*) would leave the reconstruction unchanged while dividing the L1 term by *α*; this can lead to a fitting degeneracy where as *α* → ∞, the L1 penalty vanishes and Eq. (4) no longer favors minimal-support solutions. In milder cases, the scaling degeneracy could lead to unequal operator norms, which would bias which operators are recruited since a larger-norm operator would afford more error reduction per unit of coefficient. Fixing *σ*_max_(***f***_*m*_) = 1 removes this degeneracy without restricting the model’s expressiveness, since |*c*_*m,t*_| absorbs the scale; *c*_*m,t*_ is then comparable across operators [Mudrik et al., 2024]. We minimize Eq. (4) by alternating between solving for ***C*** with ***F*** fixed, and a projected gradient step on ***F*** with ***C*** fixed, with the projection mapping each ***f***_*m*_ to the unit operator-norm sphere (Methods, §4.3).

### 2.2 dLDS identifies a 3-D space of single-mouse action dynamics

We first fit dLDS and keypoint-MoSeq to the single-mouse open-field recordings (Fig. 1a) released with keypoint-MoSeq (Weinreb et al. [2024]). This dataset, being more constrained than multi-animal social behavior, allows us to explore the properties of dLDS and the general differences between categorical dynamics from composite dynamics. In the shared 16D pose feature space, the one-step reconstruction for dLDS achieved a relative error of 2.20%, which is comparable with the 1.11% for keypoint-MoSeq (Supplementary Table S6), as keypoint-MoSeq’s AR(3) model only needs to reconstruct the 4D representational latent space and has approximately an order of magnitude more parameters (≈ 8, 000 vs. 800 for dLDS).

Keypoint-MoSeq identified 21 substantive syllables above a 0.5% usage threshold [Weinreb et al., 2024], pooled across five sessions (Fig. 1c). With the same threshold applied to the dLDS coefficients, we found that three (out of *M* = 15) operators, ***f***_14_, ***f***_2_, and ***f***_6_ accounted for 94.3%, 3.3%, and 2.3% of the coefficient variance, respectively (Fig. 1d). These operators were robustly active across a range of thresholds and regularization coefficients (Supp. Table S1b). We will illustrate how one may go about interpreting these operators in the following section.

To illustrate the differences between keypoint-MoSeq and dLDS, we consider a 7-s trajectory that keypoint-MoSeq partitioned into 9 bouts drawn from 8 distinct syllables (Fig. 1e). This bout contained a single instance of the animal turning left (Fig. 1e) over the course of 49 frames. In the syllable representation of this bout syllable 21 was active over 36 frames, transitioning to syllable 69 over 13 frames, and followed by syllable 0 which represented the slight right turn over the final 17 frames as the body reoriented into a straight position. During the transition from syllable 21 to 69, the left turn continued uninterrupted, indicating that keypoint-MoSeq split the “left turn” action into multiple syllables. In contrast to the fragmentation of actions by keypoint-MoSeq, dLDS provided a more concise, and graded description. During left turns represented by syllables 21 and 69, *c*_2,*t*_ was negative with slowly changing magnitude, *c*_14,*t*_ held near +0.7, and *c*_6,*t*_ was minimally activated. While these descriptors of the dynamics were consistent, we noted no clear changes in the dynamics description around the transition time between keypointMoSeq syllables 21 and 69 (Fig. 1f). Like keypoint-MoSeq, dLDS identified the right turn as different from the earlier left turn. However, while keypoint-MoSeq used syllable 0 for the right turn, dLDS used a single operator for both turn directions, with a negative coefficient *c*_2,*t*_ for left turn and positive *c*_2,*t*_ for the right turn (indicating a curling in the “opposite” direction, i.e., uncurling). Interestingly, the dLDS coefficients captured a change in the “uncurling” that increased over syllable 0 bout duration, which may correspond to the non-homogeneous speed of the action. The “turning right while forward” movement in dLDS is represented as a graded up *c*_14,*t*_ and a *c*_2,*t*_ first sustained then graded down to zero (Fig. 1f, bottom).

This example shows how one-hot, categorical models, exemplified here by keypoint-MoSeq, forcibly partition the continuous action manifold (Fig. 1f). In the dynamical systems setting we explored, we can view each syllable’s dynamics matrix as an isolated point on the action manifold, and any action it its vicinity is classified as that syllable, effectively splitting the continuous space of actions into discrete partitions. This means that two actions that are close together on the manifold can fall on different sides of a partition boundary and thus be assigned to different categorical syllables despite being highly similar. dLDS removes the one-hot constraint, allowing the space of possible dynamics ***A***_*t*_ to be represented using a continuous basis instead of an arbitrary discretization.

#### dLDS operators define signed action features

As each dynamical operator in dLDS represents a basis element for describing the keypoint dynamics, we next explore what the operators represent and how to interpret them. Because the operators are linear, we can analyze their eigendecompositions to characterize and understand the individual linear operators. Specifically, we computed the eigendecompositions of the three primary operators (***f***_14_, ***f***_2_, and ***f***_6_) and explored their dominant eigenvalues and eigenvectors. A dominant eigenvalue, *λ*, provides information about the general behavior of the operator. If *λ* = 1, the action mode is stable over time, *λ <* 1 attenuates the mode’s effect and |*λ*| *<* 0 creates rapid oscillations. Slower oscillations can be achieved if the top two eigenvalues share the maximum magnitude but are complex conjugates of each other. The eigenvectors corresponding to the top eigenvalue(s) provide complementary information about what configurational changes are controlled by the dominant modes.

In analyzing the dominant eigenvalues, we found that ***f***_14_ had one large dominant mode at *λ* = +1.00, with all others at |*λ*| ≤ 0.13. This sharp dropoff indicated that ***f***_14_ captured a persistent one-dimensional mode. By contrast, ***f***_2_ and ***f***_6_ each had two dominant real eigenvalues of opposite sign (−0.76, +0.74 and −0.65, +0.57, respectively), comparable in magnitude within each (Fig. 2a). This split suggests that each of ***f***_2_ and ***f***_6_ represent two, related, transient modes. To better understand if these modes were related, or if dLDS was simply multiplexing modes within the same operator, we next explored the eigenvectors associated with the top eigenvalues.

**Figure 2:**
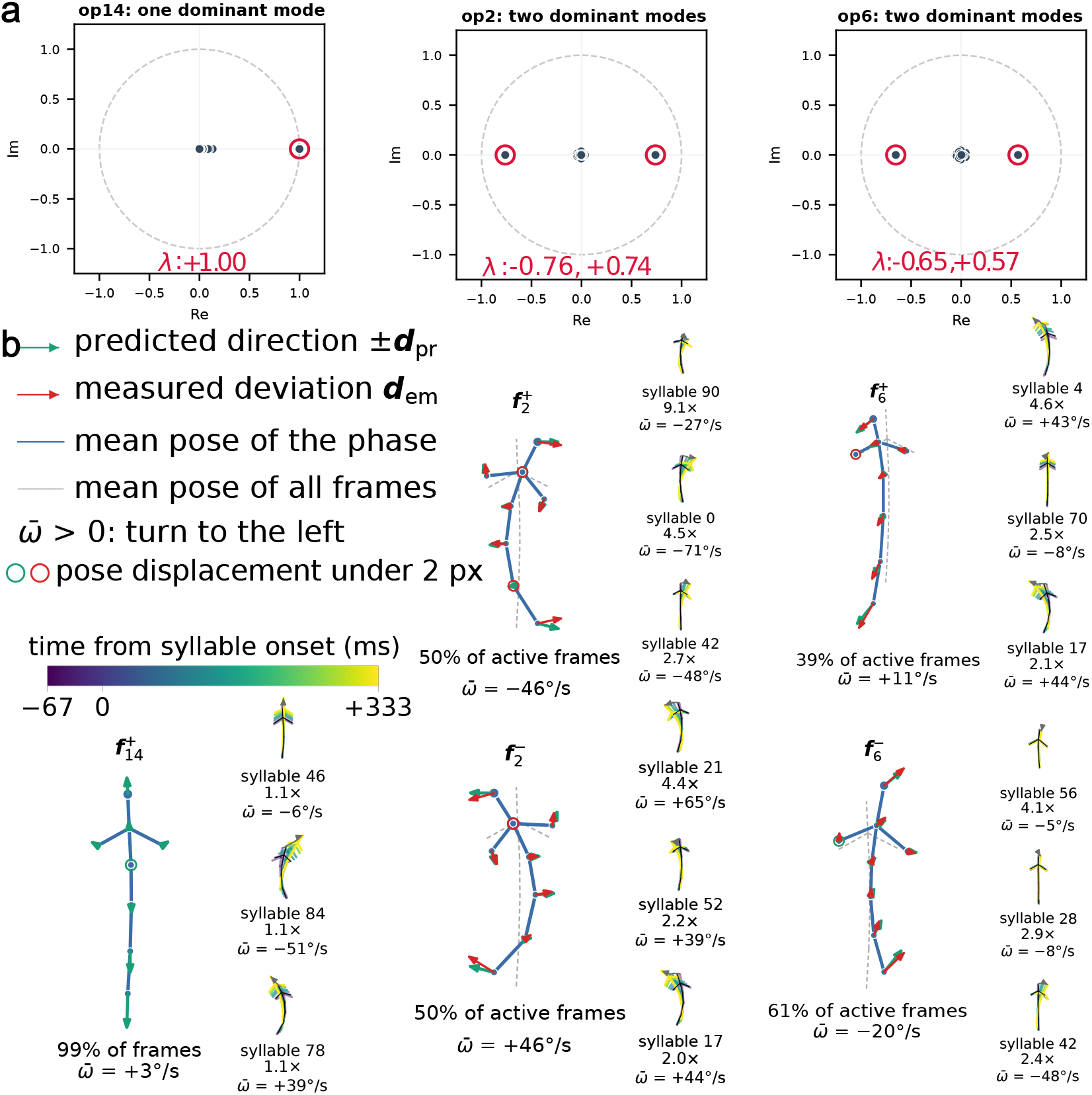
Each operator reads as one signed movement axis that recurs across several keypoint-MoSeq syllables. **a)** Eigenvalue spectra of ***f***_14_, ***f***_2_, and ***f***_6_ in the complex plane. **b)** At the left of each phase block, signed operator directions are drawn as keypoint-wise arrows set on the empirical mean pose skeleton, giving the direction and relative magnitudes within the same operator; all the skeletons in later figures follow this convention. At right are the three most co-occurring onset-aligned keypoint-MoSeq syllables drawn as in Fig. 1c, evaluated by syllable enrichment (see Methods. 4.10). The ***f***_14_ percentage is the fraction of all frames with *c*_14_ *>* 0; paired ***f***_2_ and ***f***_6_ percentages give the positive and negative shares among signed active frames (*c*_*m*_ ≠ 0). Labels report median session-level occurrence *z* and mean angular velocity 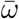.

To explore the operator eigenvectors, we visualized the keypoint displacements as predicted by each dominant eigenvector(s) as arrows pointing from each node on the mouse skeletal diagram. The diagrams thus show eight 2D vectors that start at ***f***_*k*_’s empirical mean pose keypoints (Fig. 2b) and extend out in the direction and magnitude determined by the dominant eigenvector(s). The mean pose represents the average pose that the animal takes when the given operator is active (*c*_*m*_ ≠ 0), which allows us to view the operator’s action in contexts where it is actually used. To further test the sign-specificity, we separate each operator’s empirical mean poses into two: one during times where *c*_*m*_ *>* 0 and one for *c*_*m*_ *<* 0. The arrow directions derived from the matrix agree quantitatively with the measured pose displacements, beyond a per-session circular-shift null (Supplementary Fig. S2a).

From these diagrams, we found that ***f***_14_ is always positive and corresponds to a subtle body elongation consistent with forward movement or pausing. During positive *c*_2_ epochs, ***f***_2_ represented rightward turning from a right-turn starting position. Interestingly, during the epochs where the coefficient *c*_2_ was negative, the other eigenmode of the system activated and ***f***_2_ swapped to representing a left turn from a left-turning position. ***f***_6_ had a similar opposing phases with the sign of *c*_6_, however rather than turning, the movement was more in line with a curling/translational motion.

In order to compare our eigenvector analysis with keypoint-MoSeq syllables, we estimated the co-occurrence of syllables and operator activations and identified the syllables that were most enriched for each operator (Methods 4.10). This enrichment analysis demonstrated a consistent relationship between dLDS operator activations and keypoint-MoSeq syllables. To check consistency of turning operators in particular, we computed angular velocity *ω* frame by frame from the raw keypoint trajectory, then averaged them over operator-active or syllable-labeled frames as 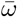 (see Methods 4.6). During frames where +***f***_2_ was active, we found that 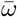 was approximately −46^*°*^ ± 14^*°*^*/*s, reinforcing the interpretation as a right turn. Similarly, the three keypoint-MoSeq syllables that were most co-active with +***f***_2_, also represented “turning right” at different degrees (−71^*°*^ ±16^*°*^*/*s, −27^*°*^ ±15^*°*^*/*s and −48^*°*^ ± 12^*°*^*/*s in Fig. 2b). The same consistency held for negative occurrences of −***f***_2_ and left turns.

This observation of multiple keypoint MoSeq syllables representing the same turn as a single dLDS operator confirms that one dLDS operator can capture the effect of several MoSeq syllables simply by changing its coefficient. This same phenomenon is observed in our 7s window (Fig. 1f) where a single turn was represented with 8 keypoint-MoSeq syllables, each with distinct angular velocity 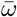. However, only two of the three dLDS operators, in various linear combinations, were needed to capture the same range of 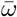 for keypoint-MoSeq syllables. In the following section we explore these many-to-many relationships across the full behavioral dataset(Fig. 3).

**Figure 3:**
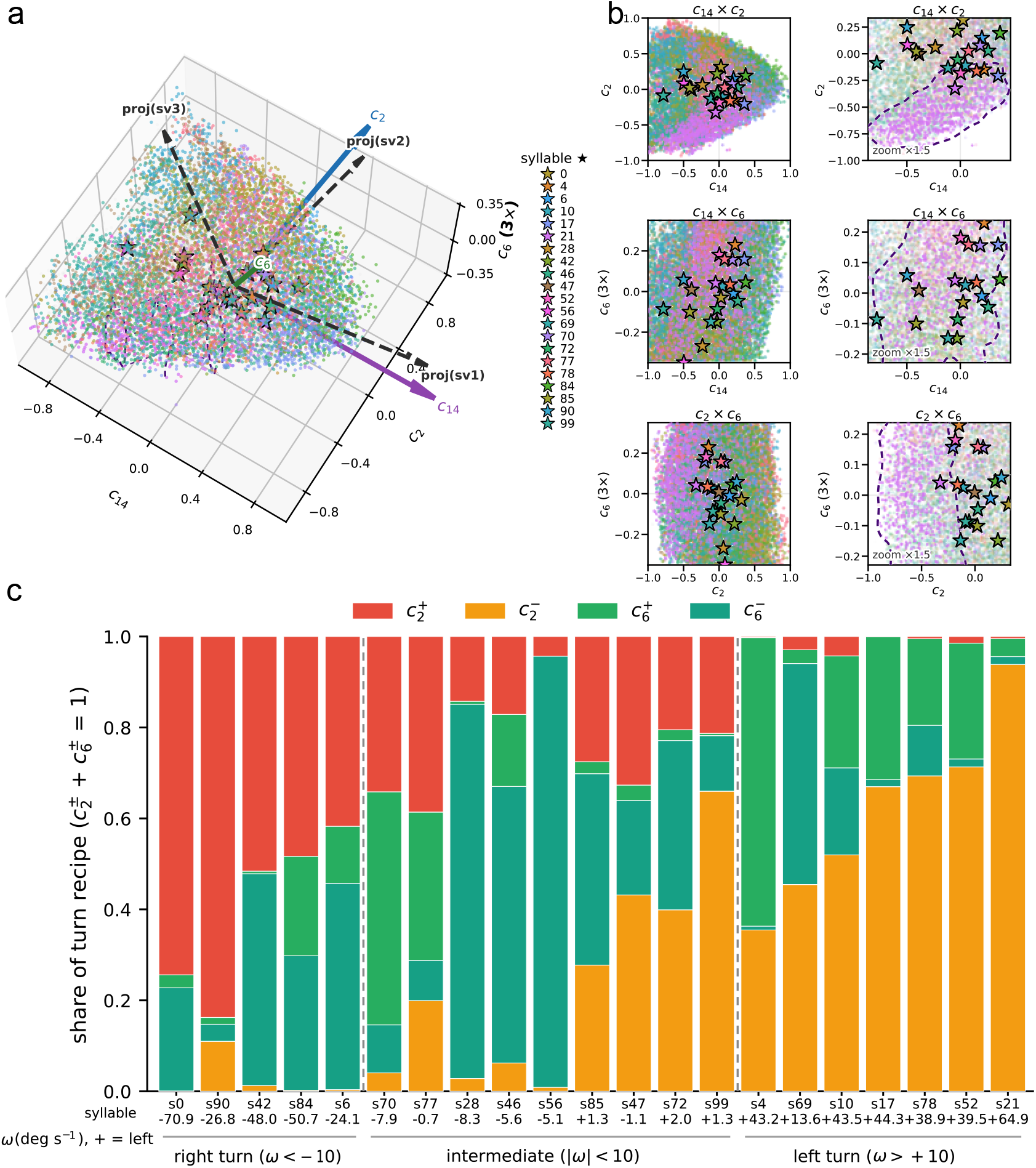
dLDS maps an action space with continuous operator coordinates while keypoint-MoSeq covers it with overlapping syllables. **a)** Single-mouse pose deviations from the dataset mean pose were projected into the orthonormal subspace spanned by the matrix-predicted directions of ***f***_14_ (forward hold), ***f***_2_ (strong turn), and ***f***_6_ (weaker turn; displayed at 3× scale). Frames are colored by keypoint-MoSeq syllable for the 21 substantive syllables and grey otherwise; stars mark syllable centroids. Dashed arrows show the top three singular vectors of the 21 × 16 matrix of syllable mean deviations, projected into the operator subspace (arrow length gives the fraction lying within it). **b)** Three 2D views of the three-dimensional operator space, column one at the same scaling, column two zoomed in 1.5× and with dashed line contours around syllable 21. **c)** Turn-composition of the 21 substantive syllables. Bars show per-syllable means of the positive and negative parts of ***c***_2_ and ***c***_6_, normalized to unit sum.

#### A shared three-dimensional action space organizes syllables

To explore the relationship between the two models’ embeddings, we analyzed how each model represents the mouse’s actions in a common same low-dimensional space. We first defined a 3-dimensional space, where each frame was represented by dLDS operators for the three most frequent operators ([*c*_2_, *c*_6_, *c*_14_]). Each point was then colored by the corresponding keypoint-MoSeq motifs corresponding to the syllable used during the same time point (Fig. 3a). To check if the 3D dLDS-based space was also a good basis for the keypoint-MoSeq representation, we constructed a 21 × 16 matrix where each of the 21 columns was the average pose coordinates for each syllable. This matrix was approximately rank-3, with the top three singular vectors accounting for 98.4% of the variance. Interestingly, the space spanned by these singular vectors was very highly aligned with the dLDS operator space, with an 88.7% overlap. (see Methods. 4.6). Specifically, the two principal angles between the two three-dimensional subspaces in 16-D were only 4.5^*°*^ and 8.0^*°*^; and the third principal angle, between the third singular vector and the operator space’s weakest axis (***f***_6_), was 45.0^*°*^ (see Methods).

While the keypoint-MoSeq syllable centroids are scattered across all three dimensions, the data cloud of operator activations does not appear to aggregate into distinct clusters (Fig. 3a). Instead, keypoint-MoSeq syllables appear to tile a continuous “manifold”, appearing as highly overlapping point clouds (Fig. 3b). From 2D slices of the 3D space (e.g., the *c*_2_, *c*_14_ space; Fig. 3b, left), it is apparent that keypoint-MoSeq’s partition-based labeling does group frames into noticeable “sub-manifold” regions, which are especially visible after zooming in (Fig 3b, right). Syllable 21, for example, covers nearly the whole range of ***c***_6_ and the negative range of ***c***_2_ from approximately − 0.75 to 0; and spans a clearly visible purple area in each projected plane.

We next sought a more quantitative description of exactly how the various linear combinations of dLDS operators map onto the syllables. We computed for each keypoint-MoSeq syllable the average activation of each dLDS operator (see Methods). Specifically, we computed the proportion of time each operator, with positive or negative signs, occurred instead of absolute magnitudes to eliminate differences caused by how long the animal spent in each syllable. We excluded ***f***_14_ from the proportion recipes, as it is simply a baseline controlling the forward/hold action.

We found that the relative proportions of positive and negative occurrences of operators ***f***_2_ and ***f***_6_ changed considerably from right to left turns (Fig. 3c). Specifically, we observed a graded decrease in 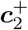 as syllables progressed from a strong right turn 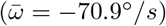 to a moderate right turn 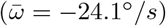. Similarly, 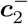 increased gradually as syllables sped up their left turns. Intermediate action syllables had comparatively balanced operator configuration (Fig. 3c), with ***c***_6_ taking a sizable and sometimes dominant share. This is because ±***f***_6_ had smaller 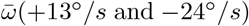 and were involved in the action where a turn starts or ends, that is, a curling or translational motion.

### 2.3 dLDS factorizes social behavior into self-, partner-, and social-actions

#### Dyadic encoding sorts operators into three types

Given the ability of our approach to compositionally characterize the action space for a single animal, we proceeded to test our hypothesis that compositional action spaces are can similarly characterize social actions across many animals. We thus trained dLDS on dyadic data, specifically to 69 sessions of the CalMS21 dataset that consisted of free interactions between mice in a resident / intruder assay. Each session’s data consisted of seven keypoints per mouse, and each frame was hand-annotated as “attack”, “investigation”, “mount”, or “other” actions (Fig. 4a). Keypoint-MoSeq (as in Weinreb et al. [2024] where it was run on the same dataset) identified 28 syllables above 0.5% usage from a 98-syllable dictionary on the resident trajectory (Fig. 4b).

**Figure 4:**
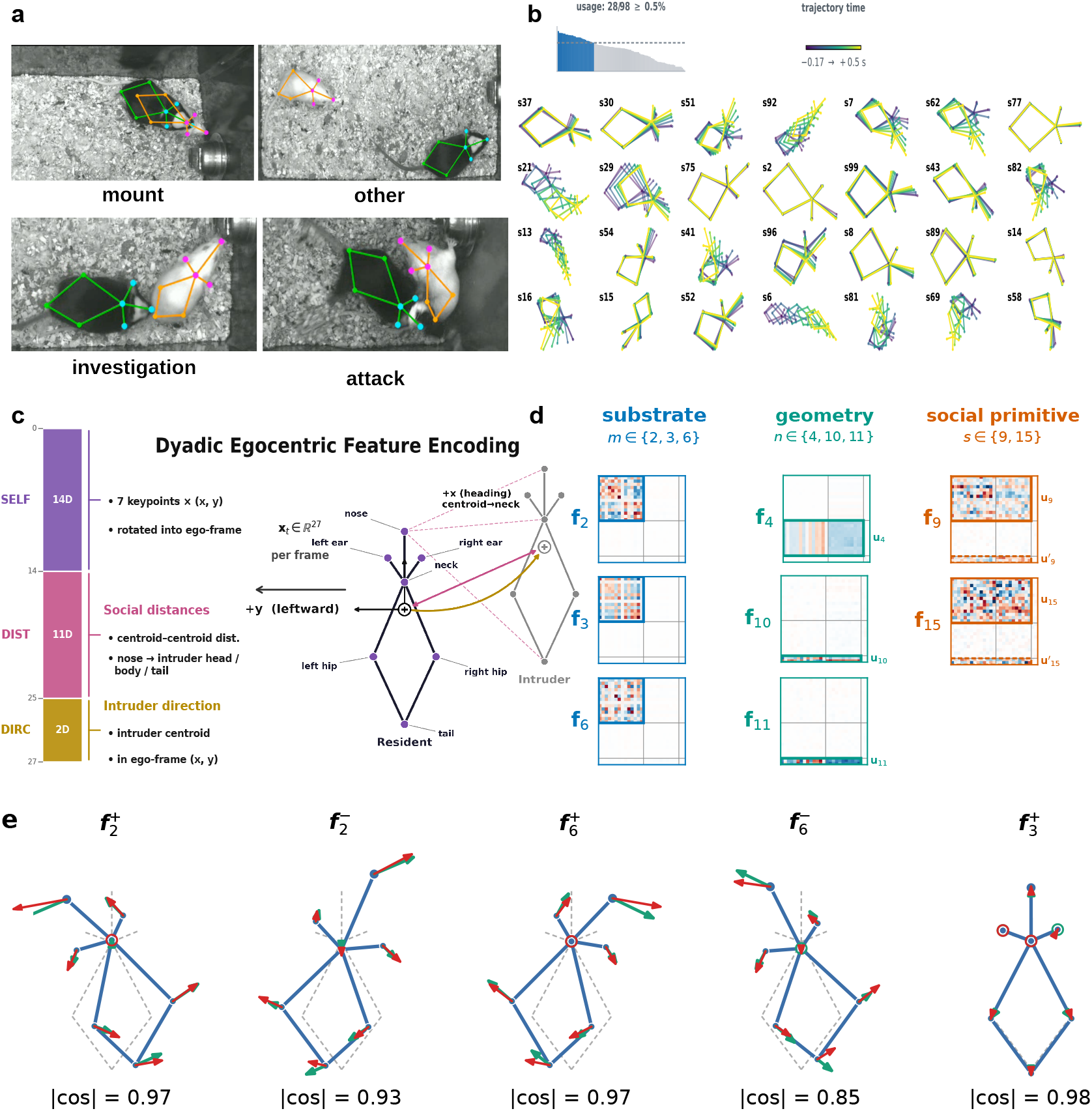
Dyadic behavior is represented by a signed mixture of self-action, intruder-geometry, and social-action operators. **a)** Representative CalMS21 frames from “attack”, “investigation”, “mount”, and “other”, with the seven resident and seven intruder keypoints overlaid. **b)** 28 dyadic keypoint-MoSeq syllables (≥ 0.5% pooled usage; *κ* = 3 × 10^4^), ordered by usage, with a long low-usage tail below. **c)** The 27-dimensional per-frame state concatenates three blocks: resident self-action (SELF), inter-animal distances (DIST), and intruder direction (DIRC). The resident centroid is the origin, +*x* the centroid-to-neck forward axis and +*y* the resident’s anatomical left. **d)** The eight interpreted operators of the learned 15-operator dictionary (Supplementary Fig. S1); the rest of each matrix is faded. Boxes mark each operator’s dominant blocks. For ***f***_9_ and ***f***_15_, solid boxes indicate resident action and dashed rows indicate intruder-direction blocks. **e)** All five phases of the three substrate operators, each drawn at its mean pose over its active frames (|*c*_*m*_| *>* 0, pixels). Green arrows show the leading eigen-mode of the SELF ←SELF block. Cosine similarity between action predictions (green arrows) and measured pose deviation (red arrows, *d*_emp_) is indicated for each operator.

To allow dLDS to learn both single-animal and social actions, we learned dynamics over a pose state space that included both the resident and intruder keypoints. Specifically, each dLDS state ***x***_*t*_ was defined as a 27-dimensional vector (Fig. 4c) concatenating three blocks of coordinates: 1) the 2-D locations of the seven keypoints defining the resident’s pose in its egocentric frame (SELF block, 14 dimensions), 2) inter-animal distances, between matching body parts, centroids, and from the resident’s nose to the intruder’s nose, neck, and tail base (DIST block, 11 dimensions), and 3) the intruder’s direction in the same egocentric frame (DIRC block, 2 dimensions). Maintaining a separation in state space between the kinematic coordinates of the two animals and their distances enables a more interpretable reading of the learned action as being specific to one animal or indicative of a social interaction and is a distinguishing feature of our approach. In contrast, keypoint-MoSeq syllables operate on a latent state ***p***_*t*_ that mixes keypoints through the principal components, making them harder to directly interpret with respect to the potential social nature of the actions.

With the 27 total pose features, each dLDS operator is a 27 × 27 matrix with a block-wise structure that indicates either animal-specific actions (interactions between keypoints within a single animal), or social interactions that include interactions between keypoints of both animals (Fig. 4d). While we allowed the model to use up to 15 total operators, there was a significant drop in operator activation after the top ten (a 9.3 × gap) and so we retained only the ten top operators for further analysis (Supplementary Fig. S1a).

Interpreting an operator in the dyadic setting depends on which features it acts on: on the resident’s own action, on the inter-animal features, or across both. Of the top ten operators, eight displayed clear block structure, with one or two output rows of the 3 × 3 block grid carrying a significantly larger share of the operator’s coupling than chance, as measured by a shuffle test (Methods 4.5, Eq. 14; solid boxes in Fig. 4d). The remaining two operators, ***f***_7_ and ***f***_8_, were not as well concentrated, so we did not assign them a class of action or intruder geometry. While not clearly in one of these classes, in their inputs and outputs, ***f***_7_ resembles ***f***_4_ and ***f***_8_ resembles ***f***_15_, with subtle functional differences, we thus treat them as secondary, less concentrated variants of ***f***_4_ and ***f***_15_ and describe them in Supp. S1.

The eight block-dominated operators fell into three categories according to which blocks were most dominant (Supplementary Fig. S1b): ***f***_2_, ***f***_3_, ***f***_6_ were “substrate operators” since they concentrated on the self-action block, ***f***_4_, ***f***_11_ and ***f***_10_, were “geometry operators” since they were concentrated on the relational blocks, and ***f***_9_, ***f***_15_ were “social primitives” since they had nontrivial weights in both blocks. This repertoire was reproducible under randomized reinitialization, with all self-action, geometry, and social-action operators recurring in 10 of 14 reinitializations.

Interestingly, the substrate operators mirrored operators found for the single animal model. By analyzing the eigendecompositions of the SELF ← SELF blocks that these operators concentrated on, we found that again there was a “sustained forward drive” 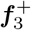, and two sign-paired turning modes with opposite left/right preference (± ***f***_2_, ± ***f***_6_; Fig. 4e, Supp Fig. S4a,b). Thus the dLDS model found a consistent set of self-actions within the larger multi-animal feature space.

#### *f*_4_ and *f*_10_*/f*_11_ represent the intruder geometry

We next analyzed the operators that act on the geometry between resident and intruder, ***f***_4_ and ***f***_10_*/****f***_11_. To develop an intuitive understanding of these operators, we made two mathematical simplifications. First, we focused on the dominant blocks: ***f***_4_ acts on the eleven inter-animal distances through its DIST rows, **M**_4_ ∈ ℝ^11*×*25^, and ***f***_11_*/****f***_10_ act on the intruder direction through the DIRC rows, **M**_11_ ∈ ℝ^2*×*25^ (Supp. Fig. S1). The contribution of an operator *k* in this simplified form can thus be written as

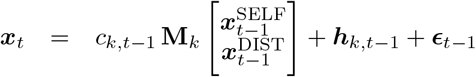

where **M**_*k*_ contains the rows of ***f***_*k*_ that act on the relevant features (DIST for ***f***_4_ and DIRC for ***f***_11_*/****f***_10_), and ***h***_*k,t−*1_ represents the minor contributions of the other co-active operators.

Second, we found that **M**_4_ and **M**_11_*/***M**_10_ were all approximately rank-one to within 1% (Supp. Fig. S3), meaning that we can analyze the output of each along the leading left singular vector ***u***_*k*_ corresponding to the largest singular value *σ*_*k*_. To isolate the contribution of each feature group, we split the right singular vector into parts that “read in” the resident’s pose and the distances, respectively: ***v***_*k*_ = [***v***_*k,s*_; ***v***_*k,d*_]. Thus, we can further write operator *k*’s contribution as:

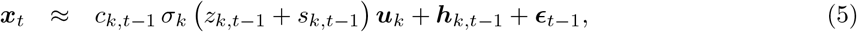

where 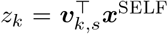 and 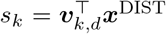 are the projections of the pose on ***v***_*k,s*_ and of the distances on ***v***_*k,d*_.

For both ***f***_4_ and ***f***_11_*/****f***_10_, we found that ***v***_*k,s*_ is highly similar to the mean egocentric pose (within 12^*°*^) and ***v***_*k,d*_ is within 10^*°*^ of the uniform vector (Supp. Fig. S3a,b). This implies that *z*_*k*_ computes the similarity between the current pose and the mean pose, while *s*_*k*_ tracks the overall separation between the animals. To validate this interpretation, we independently measured the separation of the two animals in pixels and compared this to the *s*_*k*_ values (Fig. 5a). From 200 to 770 px, *z*_*k*_ changes minimally with separation (bin medians ≤ 12%) while *s*_*k*_ grows linearly (*r* = 0.92; Fig. 5a). Thus, *z*_*k*_ + *s*_*k*_ primarily tracks the separation and is positive for over 99.5% of the frames in which ***f***_4_ and ***f***_11_*/****f***_10_ are active (Supp. Table S2), meaning that the sign of *c*_*k*_ alone sets the output direction along ***u***_*k*_.

**Figure 5:**
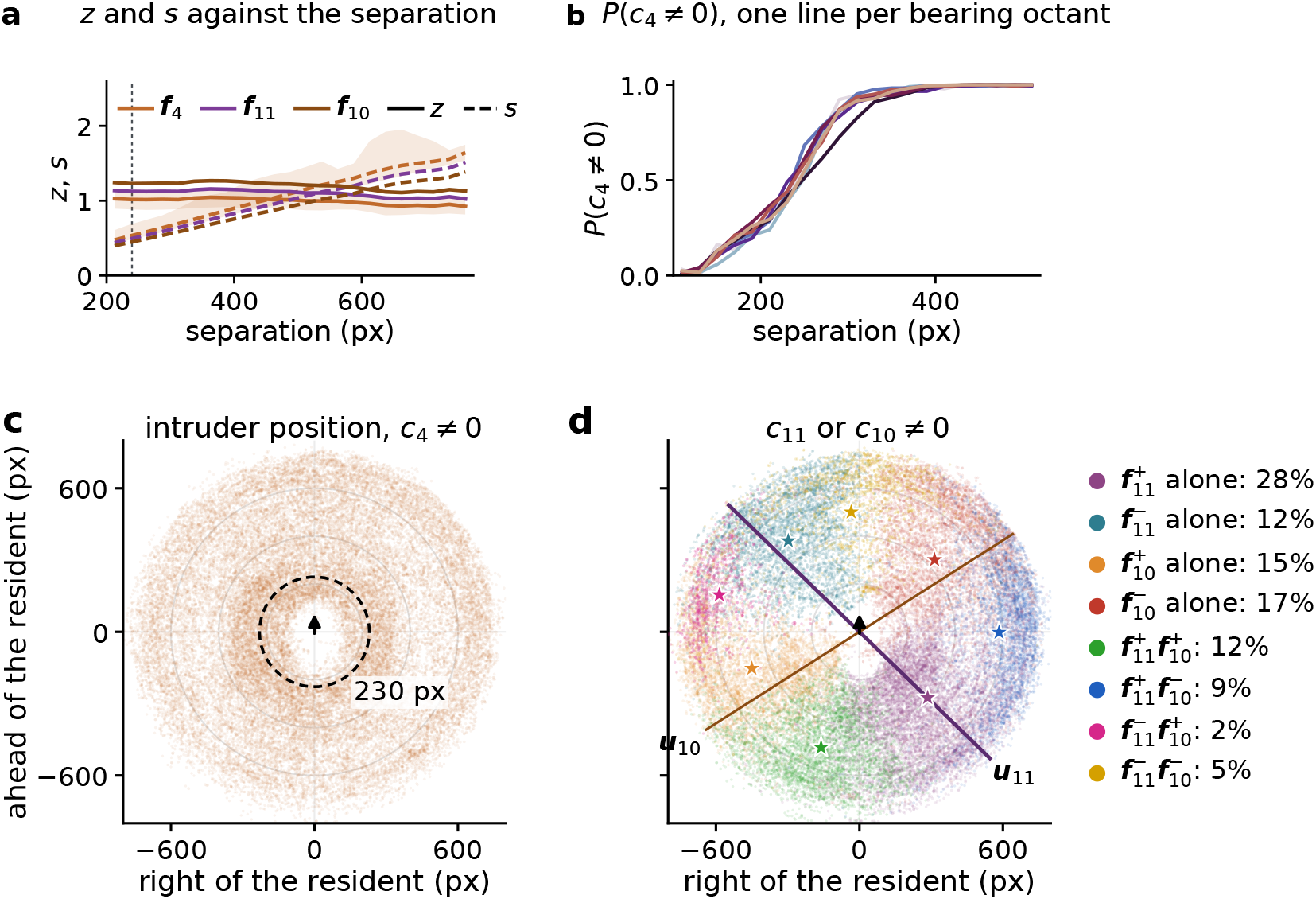
Interpreting the geometry operators. **a)**The two scalars (*z* and *s*) of ***f***_4_, ***f***_11_, ***f***_10_ as a function of average separation in 25 px bins. Lines are the median separation and shading indicates the 25th to 75th percentile range. **b)** *P* (*c*_4_ ≠ 0) along each bearing direction as a function of the mean inter-animal distance. **c**,**d)** Intruder positions in the resident’s egocentric frame with *c*_4_ ≠ 0 (left; *n* = 256, 360 points) and either ***f***_11_ or ***f***_10_ active (right; *n* = 165, 671 points).

We found that ***f***_4_ influenced the separation between resident and intruder independent of the intruder’s bearing. Surprisingly, ***f***_4_’s activation *c*_4_ signaled when the animals were apart vs. when they were close, providing a natural “proximity gate” for determining social behavior. When the proximity gate is active, the animals are distant and the inter-animal geometry is the dominant component of dynamics in the system. Alternatively, when the proximity gate is inactive, other social operators (discussed below) become active. Moreover, when *c*_4_ ≠ 0, the output of ***f***_4_ predicted the mean normalized distance of the next frame (*r* = 0.95, *n* = 254, 576 frames, Supp. Fig. S3c, left). This distance was projected onto all the outputs uniformly, due to 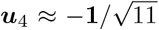 (Supp. Fig. S3b), effectively capturing the inter-animal distance dynamics. This interpretation is reinforced by ***f***_4_’s activation only depending on the separation, and not the intruder’s bearing, with increasing activation probability with increased distance (Fig. 5b), being essentially silent within ≈ 100 px (half body length), and only 10% activation within ≈ 230 px (1.2× body length of each other (Fig. 5c). We thus call *c*_4_ = 0 the *proximity gate*; frames with *c*_4_ = 0 are *near* frames, and the rest (*c*_4_ ≠ 0) are *far* frames. We found that the gate represented an meaningful social proximity, with 94% of human-annotated social behavior frames being near frames and 95% of the far frames containing “other” behaviors (Supp. Fig. S3d). Thus, *c*_4_ = 0 marks the edge of a social contact zone that the model discovered without knowledge of the social behavior labels.

Complementing ***f***_4_’s distance information, ***f***_11_ and ***f***_10_ track the bearing of the intruder relative to the resident’s heading. Together with ***f***_4_, they represent *where* the intruder is while the animals are apart. The signs of *c*_11_ and *c*_10_ give the directions of the axes on which the intruder lies, behind-right/behind-left (+) or ahead-left/ahead-right (−), and the scale |*c*_*k*_*σ*_*k*_(*z*_*k*_ + *s*_*k*_) | gives its position along 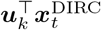 (*r* = 0.86 with k=11 and *r* = 0.78 with k =10, Supp. Fig. S3c, middle, right). Furthermore ***f***_11_ and ***f***_10_ partition the angular location space. When only one of ***f***_11_ and ***f***_10_ are active, the intruder is almost always in the direction of the active phase (99.2% and 97.6% of frames for ***f***_11_ and ***f***_10_, respectively). In the remaining frames, the combined sign combination indicates which quadrant the intruder is oriented in (93% of co-active frames; Fig. 5d). Together, the two signed operators ***f***_11_ and ***f***_10_ represent all possible intruder bearings as a point on an eight-sector colored compass (Fig. 5d).

#### *f*_9_ and *f*_15_ represent distinct, close-contact social interactions

We found that the ***f***_9_ and ***f***_15_ operators tended to occur within the “near zone” and were strongly associated with “mount” and “attack” labels, respectively. The association of 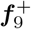 with “mount” (5.7% of frames) and 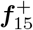 with “attack” (2.8% of frames) held over all frames as measured by the enrichment *E* = *P* (label | phase active) */ P* (label) which equals 1 when the label and phase are unrelated (*p <* 0.005, per-session circular-shift test; see Methods 4.10).

Specifically, we found that “mount” is enriched 2.94× during 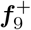, and “attack” is enriched 1.79× during 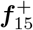. A mathematical intuition for these operators was developed using the same approach as for ***f***_4_, ***f***_10_, and ***f***_11_, noting that ***f***_9_ and ***f***_15_ were primarily concentrated in the rows for the resident’s action (SELF rows) and the intruder’s position (DIRC rows). Writing the constrained dynamics as before yields

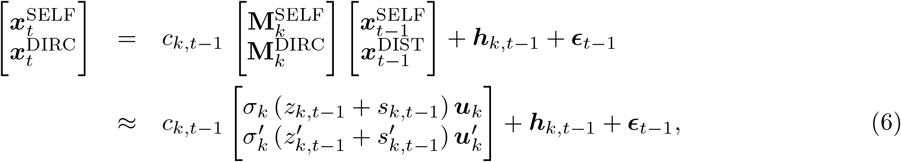

with *z*_*k*_, *s*_*k*_, 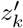and 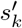 are again the projection onto each row block’s leading right singular vector. As before, a rank-one approximation captured 98% of the output energy in both row blocks for ***f***_9_, and 75% (SELF) and 83% (DIRC) for ***f***_15_. We therefore focus on the readout axes ***u***_*k*_ and 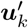 for each *k* = 9, 15. The scalars projections *z*_*k*_ + *s*_*k*_ for of ***f***_9_ and ***f***_15_ were positive over each operator’s active frames (Supp. Table S2), indicating again that the sign of *c*_*k*_ dictated relative orientation (Fig. 6a,c, left). As before, we aligned the output singular vectors with the pose positions that resulted in positive contributions to the state to inspect and validate the operator actions (see Supp. S5).

**Figure 6:**
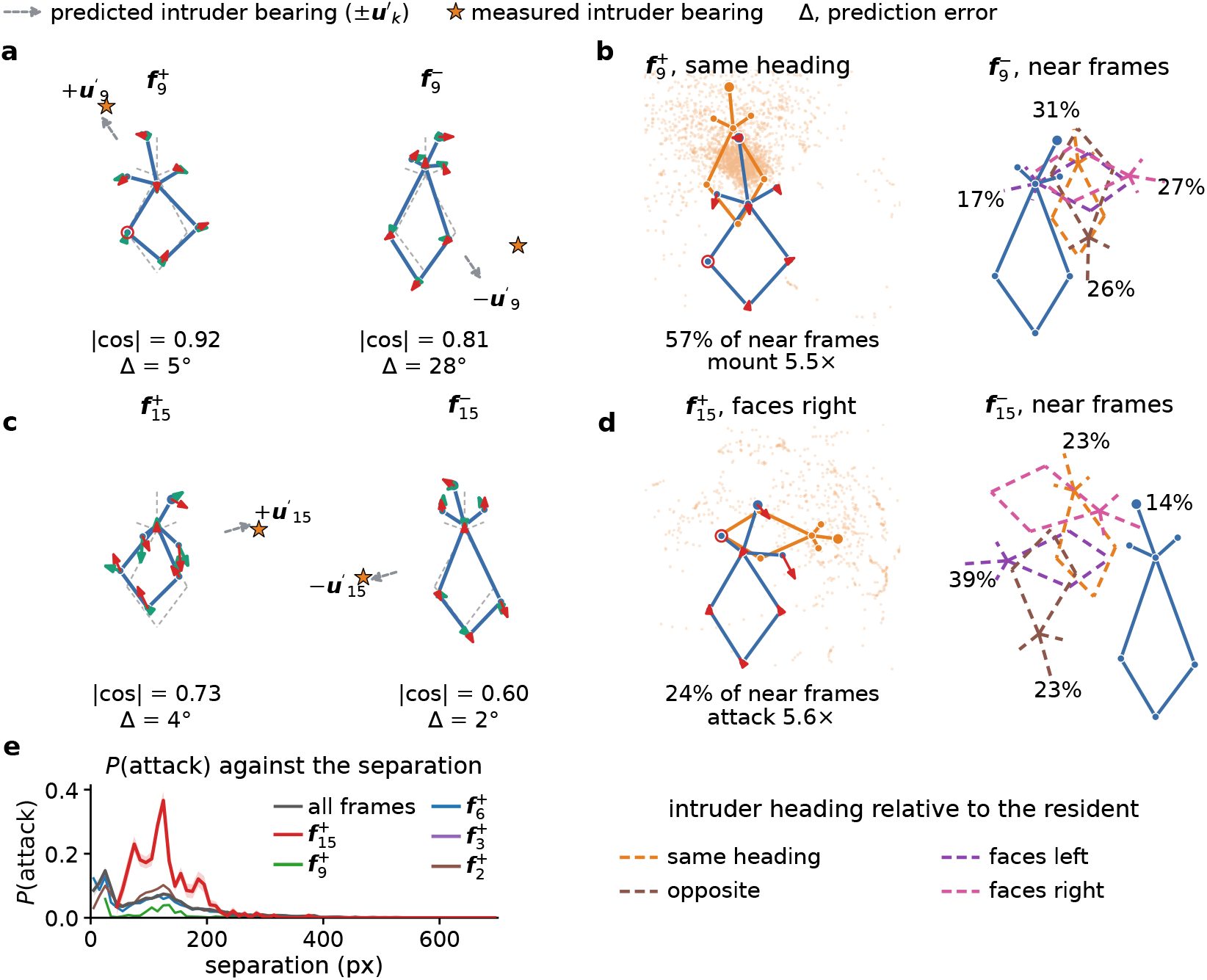
Interpreting the action and relative heading of social operators. **a)** SELF- and DIRC-row readouts of 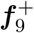 (left) and 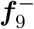 (right) for frames with *c*_*k*_ *>* 0 and *c*_*k*_ *<* 0, showing predicted action ±***u***_*k*_ (green arrows), measured pose deviation *d*_*emp*_ (red arrows), and intruder directions. **b)** Inter-animal dominant configurations for 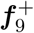 and 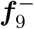. Relative heading is the angle between the intruder’s (orange) and the resident’s (blue) body axes, measured from the centroid to the neck, with intruder positions shown as dotted orange points. Left, 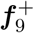 resident-centric frames within the same-heading group (relative heading *<* 45^*°*^, 57% of near frames), the “mount” enrichment within this group. Right, 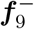 resident-centric near frames, coupled with the intruder’s four dominant relative-heading groups (dashed). Percentages indicate relative number of frames per group. **c**,**d)** The same as (a,b) but for ±***f***_15_ and attack enrichment. **e)** probability of “attack” against separation (centroid distance, 25 px bins), over all frames and social operator active frames with *c*_*k*_ *>* 0. Shaded bands are the binomial s.e.

Kinematically, 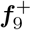 slightly shifts the resident pose to the left, with its trunk folded and neck retracted. When paired with the intruder bearing, which was in the front-left both as predicted by the model (+34^*°*^) and measured in the video (Fig. 6a), we deduced that 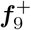 is a trunk compression *towards* the intruder. 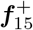 captures a large opposing swing of the resident’s head and trunk, its nose moves to the right and trunk moves to the left, with its ears folded back, neck extended and the intruder on its right, near −74^*°*^. 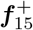 ‘s measured deviation from the global mean pose is 1.6× that of 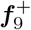 (45 vs. 28 px), with the resident’s coil showing stronger trunk compression than 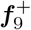 (Fig. 6c), making 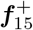 ‘s a more dramatic action than 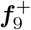.

We next explore the bearing of the intruder relative to the resident. For 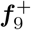, the dominant configuration has a shared heading between the resident and intruder, with the intruder located directly in front of the resident (Fig. 6b). When we sorted frames by the angle between the two animals’ trunk axes (grouped into four categories: same, left, right, and opposite) the same-heading category contained 57% of the frames, more than twice the random chance level. Within same-heading frames, the intruder is consistently located directly ahead of the resident (*R* = 0.90), with the resident’s head located over the intruder’s rump (Fig. 6b). This configuration was characteristic of mount, with a 5.5× mount enrichment over all near-zone frames. In the other three categories, the distribution of the intruder location is more dispersed (*R* = 0.67 to 0.74; Supp. Fig. S4c). The combination of the kinematics and pose statistics point to 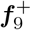 as a mounting action. Indeed, within the proximity gate *c*_4_ = 0, human-annotated mount frames were 3.6 more frequent in 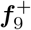 frames than in all near-zone frames.

For 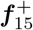, frames inside the proximity gate with *c*_15_ *>* 0 are enriched for another social behavior: “attack” (3.7× more prevalent as in all near frames). At inter-animal separations between 50 px and 200 px, which account for 90% of attacks, “attack” is 2.5 × −4.1× as frequent on 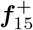 frames as on all frames, as compared to the positive phases of other operators that are *<* 1.35× (Fig. 6e). When we sorted 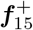 frames into the same four heading categories as we did for 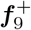 (Supp.Fig. S4d), we found no dominant category (20% to 36% each, close to chance), indicating that 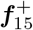 represented an orientation-agnostic behavior, which attack is (Segalin et al. [2021], Sun et al. [2021]), rather than the orientation-specific mount behavior that 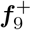 represented (Fig. 6d, left).

Interestingly, with negative coefficients, ***f***_9_ and ***f***_15_ represent different social dynamics. For one, 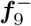 stretches the resident’s body and orients rightward (− 25^*°*^ against +12^*°*^)(Fig. 6a, right), with the intruder behind the resident to the right (− 117^*°*^, 71% of frames). Like 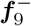, 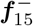 stretches the residents body as well and orients it to the opposing side, to the left (+15^*°*^ against − 30^*°*^) with the intruder behind on its left (+104^*°*^, 89%) (Fig. 6c, right). In contrast to the positive-phase, neither 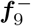 or 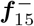 within the near zone had a dominant intruder heading (largest category 31% for 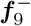 and 39% for 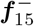; *R* = 0.14 and 0.31; Fig. 6b,d,right), nor did they have a social behavior enrichment. 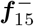 shows no attack enrichment (0.26×) and leans toward frames labeled “other” (1.80×, *p* = 0.002), while 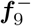 is depleted on “mount” (0.32×, *p* = 0.001) and only weakly enriched for “investigation” (1.28×).

#### Co-active dLDS operators represent rich social behaviors

In the previous section we analyzed individual operators, showing that the learned dynamics are interpretable and intuitive relative to the human-annotated behaviors. However, combinations of operator activations, and their temporal dynamics, provide a more complete view of social interactions at each moment. Here we explore these inter-operator dynamics and show how a handful of activations combine into intuitive and detailed representations of social behaviors, whose descriptive power we test by decoding expert annotations.

We start with the temporal interactions between ***f***_9_ and ***f***_4_, which emphasize how 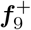 is uniquely associated with mounting. We first note that 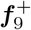 activation is strongly negatively correlated with |*c*_4_| near the onset and offset of mount bouts. By aligning *c*_9_ and |*c*_4_| to the labeled onset and offset of the mount bouts (*n* = 259), we found that the proximity gate deactivated right before bout onsets (median lead of 1.07 s) for 94% of bouts and stayed closed throughout the bout. *c*_9_ activation consistently followed ***f***_4_ silencing (median lag of 0.40 s) at mount onsets, and remained high through the bout (Fig. 7a). At offset this order reversed.

**Figure 7:**
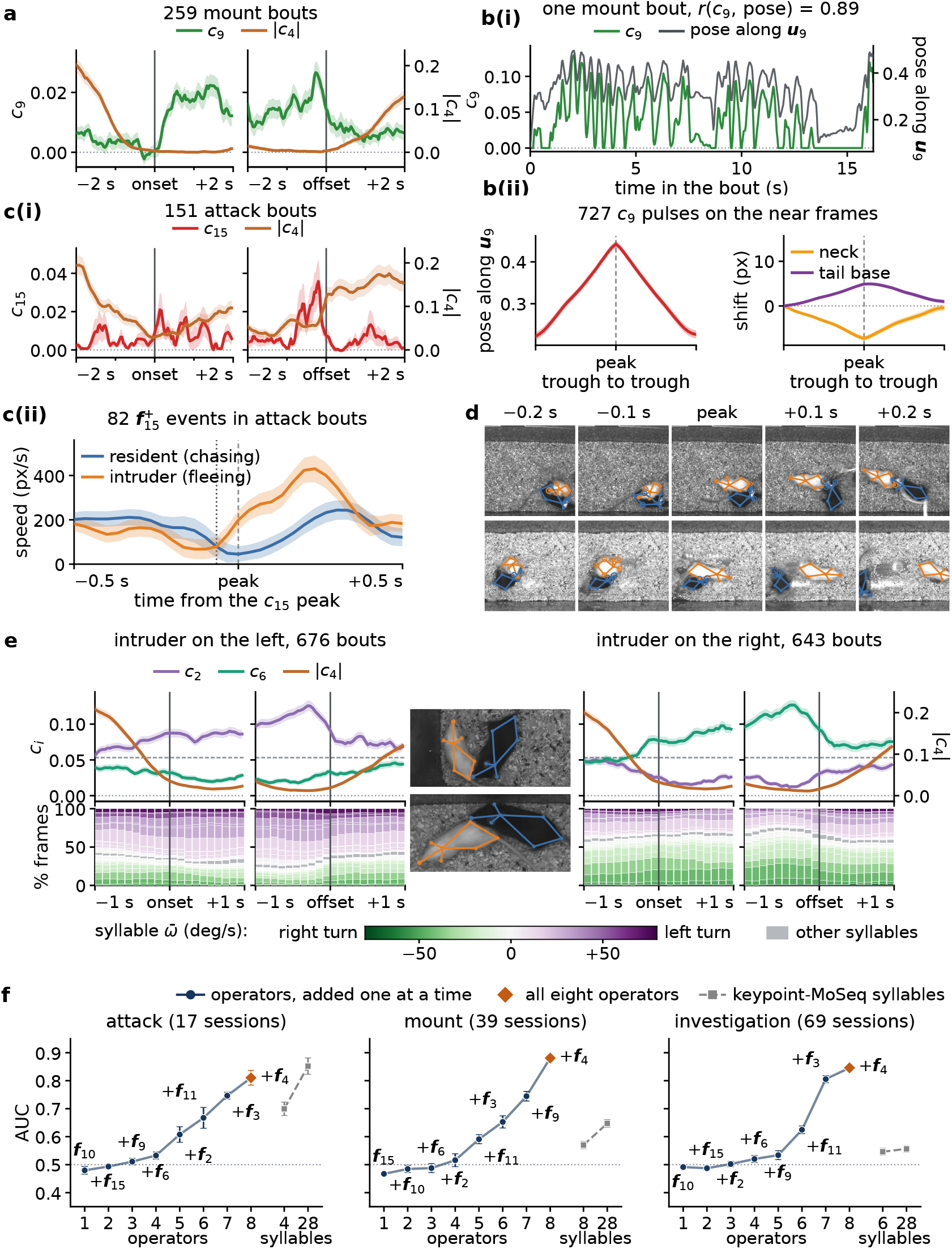
Temporal dynamics and co-activation of operators. **a)** *c*_9_ and |*c*_4_| (right axis) around the onset and offset of 259 mount bouts. Shading represents s.e.m. across bouts. **b)** Top: example 16-s mount bout dynamics. Bottom left: mean pose along ***u***_9_ aligned to the peak (peak ≥ 0.05, n=727); higher means larger body folds. Bottom Right: shift of the neck and tail base along the body axis, aligned to the pulse peaks. Shading is s.e.m. across pulses. **c)** Top: as in **a** for *c*_15_ and attack bouts. Bottom: resident and intruder speed dynamics for 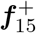 attack events (n=82), aligned at the peak *c*_15_ values. **d)** Two representative 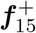 events. **e)** Investigation bouts split by the which side the intruder is predominantly in. Top, *c*_2_ and *c*_6_, and |*c*_4_| (right axis). Shading is s.e.m. across bouts. Bottom, share of frames per keypoint-MoSeq syllable in 0.1 s bins, stacked by mean angular velocity. **f)** One-vs-rest logistic regression decoding (see methods) against the number of units, iteratively adding the operator that yields the lowest AUC improvement. Error bars are bootstrap s.e. across sessions (200 resamples).

To explore the structure of a typical mount, we further examined the evolution *c*_9_ during mount bouts. In a representative 16 s bout (Fig. 7b(i), *c*_9_ rises and falls in brief pulses (see Methods. 4.8), closely tracking the pose projection onto ***u***_9_ (*r* = 0.89). To examine the resident’s kinematics within a single pulse, we averaged its body fold and the shifts of its neck and tail base along its body axis across the 727 pulses, each aligned at its *c*_9_ peak and time-normalized between its two flanking troughs (median duration 0.60 s; Fig. 7b(ii)). Averaged across these pulses (Fig. 7b(ii), left), the resident’s body fold deepened and relaxed with *c*_9_ (96% of pulses), while its neck drew back and its tail base came forward toward the peak and moved back by the next trough (75% and 71%), dynamics consistent with “mount”.

In a similar analysis for ***f***_15_ and ***f***_4_, we found that ***f***_4_ fell silent later (median lead of 0.47 s) and less completely than that for mount (88% of attack onsets fully closed). Across attacks, *c*_15_ was highest just after onset (+0.13 s) and just before offset (−0.20 s), making it sparse and tuned to attack bout boundaries (Fig. 7c(i)). To gain a more detailed understanding the inter-animal dynamics described by 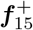, we aligned the 82 events inside attack bouts at their *c*_15_ peak, and averaged the resident’s chasing speed and the intruder’s fleeing speed along the line connecting their centroids (Fig. 7c(ii)). Combined with the kinematics of ***f***_15_, we found that 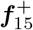 describes a rapid succession of chasing by the resident, and fleeing by the intruder. Specifically, the resident chased the fleeing intruder at a high speed (121–211 against 68–182 px/s) until 0.07 s before the *c*_15_ peak, when the two animals were closest. After that moment, the resident slowed, while the intruder sped up to ≈ 2.5× its mean fleeing speed (430 px/s at 0.23 s after the peak, against 171 px/s over all attack frames) and ran away from the resident (85% of the events). Individual representative attack events are consistent with this analysis (Fig. 7d).

Interestingly, while “investigation” was the most frequently labeled social behavior (29% of all frames) we did not find that it was associated with a single operator. This finding was consistent across 14 dictionaries fit with different random seeds, matching the lack of investigation-specific motifs in keypoint-MoSeq. However, since “investigation” bouts primarily involved the resident turning its nose toward the intruder on either side, (see bout examples in Fig. 7e, middle), we found that “investigation” labels were better predicted by the *combination* of self-action (***f***_2_ and ***f***_6_; Fig. 7e, top) and social distance (***f***_4_) operators. As with other social bouts, the proximity gate marked bout onset (median lead of 0.27 s) and offset (median lag of 0.67 s). Keypoint-MoSeq likewise had no investigation syllable(s), (best enrichment only 2.3× across 1% of investigation frames and used all 25 syllables to cover the bulk (95%) of investigation frames that dLDS captured with only the substrate operators (Methods 4.6; Fig. 7e, bottom). This result is a prime example of the need for compositionality in behavioral modeling in order to accurately describe behaviors relevant to the laboratory.

Finally, we turn to a decoding analysis to understand how the composition of operators impact decodability of human annotations. Overall, the eight dLDS operators were sufficient to decode the three social behaviors more accurately than the behavior-specific keypoint-MoSeq syllables, as measured by task-frame AUC (Fig. 7f; see Methods 4.9). When including all 28 keypoint-MoSeq syllables, the eight dLDS operators decoded “investigation” and “mount” far better with respect to the ROC curve AUC (0.85 and 0.88 for dLDS vs. 0.56 and 0.65 for keypoint-MoSeq) and decoded “attack” comparably (0.81 against 0.85). Moreover, analysis of how the decoders leveraged each operator corroborated our deduced interpretations of them. Specifically, beyond the hold (***f***_3_) and proximity (***f***_4_) operators that were shared by all behaviors, “mount” was most improved by including ***f***_9_ (+0.09 AUC increase), and “investigation” from the substrate and geometry operators: ***f***_3_ (+0.18), ***f***_11_ (+0.09) and ***f***_4_ (+0.04). Interestingly, when considering the “attack” label, the dLDS operators and the keypoint-MoSeq syllables decoded attack comparably despite keypoint-MoSeq being restricted to only the resident’s dynamics, suggesting that “attack” had a more distinct resident action set than the other social behaviors.

## 3 Discussion

In this work we developed a dLDS framework for learning compositional dynamics that represent interpretable actions based on pose information. Our framework learns, in an unsupervised way, the configurations and time courses of behavior on separate axes. The compositional nature of the model means that individual learned operators each represent individual actions that can be combined to form more complex actions. This composition is more flexible than previous relaxations of the one-regime-at-a-time commitment, which let a single operator drift smoothly across windows [Harris et al., 2021] or combined drift with switching [Lee et al., 2023]. This flexibility was crucial to capturing more nuanced social behaviors, which can contain both independent per-animal actions as well as coordinated actions across multiple animals. In our specific case of two interacting mice, we found that the configural axis and kinematic features of the learned operators were directly relatable to individual actions and social geometry. The temporal evolution of the dynamics activations further enabled us to interpret which geometric features were important to decoding social behavior. The structure of our model serves to recast behavior as a composition of reusable dynamical components, consistent with longstanding ideas of hierarchical behavioral organization [Lashley, 1951, Baerends, 1976, Dawkins, 1976], and providing a complementary view to modern syllable-based descriptions built around discrete motifs [Wiltschko et al., 2015, Markowitz et al., 2018, Weinreb et al., 2024].

While our end goal was modeling social behavior, we found that analyzing single-mouse behaviors illustrated how compositional dynamics can provide an efficient representation of action. Syllable-based models, by design, quantize potentially continuously varying actions into several motifs, such as turns at different angular velocities. In dLDS, these variations share one configural operator, and their timing and magnitude instead show up as modulations to the coefficient trajectory. Interestingly, the self-action operator structure was independently and robustly learned in both the single-mouse and two-mouse settings, indicating a potential universal description of mouse behavior (Supp. S7).

Interpretability plays a central role in behavior analysis. For this reason the sparsity in dLDS was key to ensuring we could connect the dynamics operators learned to the specific behavior. Keypoint-MoSeq is the extreme case of a single operator being on at a given time, while dLDS relaxes this assumption slightly to allow some (but not all) operators to be active, thereby permitting compositional representations. By varying the model parameters, in particular the sparsity parameter *λ* we can titrate the data fidelity vs. sparsity, allowing us to find a balance between these two requirements (Supp. S6).

One practical benefit of our framework is its scalability. We found that dLDS was faster and more memory efficient to train (see supplement S3 S4). This scalability means that we expect that adding additional animals, or increasing the number of keypoints per animal should not incur significant additional computation. Practically we also found that the pre-processing stages were key to determining the timescale of the recovered behavior. In our methodology we interpolated low-confidence keypoints and smoothed them (similar to more recent methods e.g., lightningpose [Aharon et al., 2026, Biderman et al., 2024] that incorporate smoothing directly), whereas keypoint-MoSeq corrects noise natively in its latent dynamical model. This distinction is apparent during attack bouts. Keypoint-MoSeq separates attack-associated motion into syllables with different speeds, whereas the attack-associated dLDS operator does not encode a fixed velocity regime. Instead, it captures short transition events spanning chase, closest approach, and separation. Thus, smoothing may trade sensitivity to the fastest kinematic transitions for a coarser interaction-level representation.

While human annotations provide an important reference for evaluating behavioral models, manual annotation can be coarse (combining many pose/action combinations) and incomplete (missed bouts, etc.) [Karigo and Charles, 2025]. For example, in the case of “mount”, human labels and operators capture structure at fundamentally different scales. Human labels span the mount event, from the initial climbing onto the intruder to the dismount. The operators captured the more nuanced approach and in-bout dynamics, producing a more detailed account of the resident’s in-mount motion, which can be indicative of other information such as vigor. Another important example is the attack. Human labels mark attack as one continuous bout, however the most directly related operator for attack was ***f***_15_, which was active only during a few brief transitions at the start and end of the bout. This difference meant that while ***f***_15_ was clearly indicative of attack, and quantitatively enriched in its use during attack, it only improved attack-decoding AUC slightly (Fig. 7e) since the decoding needed to represent the persistant, continuous human labels. These examples show that more fine-grained information can be crucial to understanding neural activity, e.g., vigor [Shadmehr and Ahmed, 2020, Dudman and Krakauer, 2016]. Given that dLDS, the model that forms the basis of our work, has already been used as a model of neural activity [Mudrik et al., 2024, Yezerets et al., 2026, Mudrik et al., 2025, Chen et al., 2024], a natural next step would be merge these frameworks to find joint models of motor and neural actions.

Several extensions could further separate configuration from dynamics and enrich the representation of social interactions. For one, dLDS can also include a linear latent state, similar to keypoint-MoSeq. Such a data-driven observation model could absorb recurring pose structure and observation noise, so that the operators describe the transformations around those configurations more selectively, at the cost of direct interpretability of model parameters. Expanding the dyadic feature space with more inter-animal features, and reducing the influence of long non-interaction periods, could let the model resolve a broader range of interaction-specific dynamics. With simultaneous neural recordings, the same framework could then decompose behavior and neural activity side by side and test their alignment directly.

## 4 Methods

### 4.1 Dataset and preprocessing

We fit dLDS to two keypoint datasets: 1) a single-animal dataset and 2) a full two-animal dyadic social dataset. The data represented keypoints extracted from top-down videos as described in the source publications [Segalin et al., 2021, Weinreb et al., 2024].

#### Single-mouse open-field dataset (keypoint-MoSeq)

We use the single-mouse open-field recordings analyzed in and released with keypoint-MoSeq [Weinreb et al., 2024]. This dataset consists of ten DeepLabCut-tracked sessions of individual mice totaling ~ 644,000 frames at 30 Hz. Each session contains a top-down, single-camera video recording of one mouse exploring an open-field arena. For this dataset, we retained a subset of eight body-part keypoints, dropping only the tail base to be consistent between the single and two mouse datasets.

### CalMS21 dyadic dataset

For the social mouse data we use the Caltech Mouse Social Interactions dataset [Sun et al., 2021]. CalMS21 is a dyadic mouse social benchmark. The social interactions follow the resident–intruder paradigm, wherein a naive intruder is introduced into a resident’s home cage. For this data, the pair is filmed from above at 30 Hz under infrared illumination. The CalMS21 authors used MARS (Multi-Animal Real-time Scorer) [Segalin et al., 2021] to track seven anatomical keypoints per animal: nose, left and right ear, neck, left and right hip, and tail base. Frame-level annotations cover four mutually exclusive categories: “attack”, “investigation”, “mount”, and “other” (Fig. 4a). Other here represents background or non-social activity. We used the standardized labeled training-testing split of CalMS21, in which 70 sessions totaling approximately 508,000 frames are the training data and the rest are for testing. We exclude one session whose length is only 96 frames (~ 3 s, well below the median of 5,568 frames) and retain all other 69 training sessions. We use the pose trajectories as model inputs, consistent with the unsupervised nature of dLDS, and use the labels only for post-hoc evaluation of the learned model.

#### Preprocessing

We curated the data with a minimal set of pre-processing steps to remove jitter and uncertain data-points. Specifically, we isolate keypoints and sessions that have clear errors, replacing them with locally interpolated values.

##### Low-confidence keypoints

In both datasets, we remove any keypoint on any frame with detection confidence below 0.5. We replace any removed keypoint by linearly interpolating that keypoint’s time-series between the nearest reliable values. In the single-mouse data, as in keypoint-MoSeq, a keypoint is also treated as low-confidence when its distance from the per-frame median of all keypoints exceeds its session median by more than six median absolute deviations. We thus also remove these keypoints and similarly replace them using the same linear interpolation procedure.

##### Session-wise and full-pose failures

To prevent erroneous sessions from biasing our analyses, we removed any session that did not pass basic quality tests. For the single-mouse session, the quality test checked if the nose or “spine4” keypoint is consistently a low-confidence keypoint measured by either ≥ 1 s continuous low-confidence keypoints or low confidence on ≥ 5% of all frames. Five out of ten sessions (≈ 370, 000 frames) pass these criteria and were used. For the CalMS21 dataset, we treated any frame in which ≥ 60% of the 14 keypoints are low-confidence as a “full-pose failure” (0.9% of frames, mostly isolated frames). Since full-pose failures were typically isolated frames, we likewise interpolated their keypoints from nearby frames.

##### Smoothing

The CalMS21 keypoints exhibited jitter even when the animals were still, indicating small errors in the keypoint detection. We therefore smoothed the keypoint time-series by temporally filtering the time-series with a Gaussian kernel (*σ* = 1 frame) to suppress this tracking jitter (§4.2). To retain consistency with the Keypoint-MoSeq paper, the single-mouse data keypoints were not smoothed.

### 4.2 Egocentric interaction feature encoding

We encode each mouse in a resident-centered egocentric frame (Fig. 4c). The self-action block (SELF) is thus the same in both datasets and represents the full set of keypoints for the single-mouse data, and the resident’s keypoints in the dyadic data. For the dyadic data we extend the state by adding inter-animal distances (DIST) and intruder ego-direction (DIRC), giving the 27-dimensional raw feature timeseries ***y***_*t*_. As the single animal data organization is more straightforward (and a subset of the dyadic data), we focus here on the dyadic state description.

#### Resident-centered coordinate frame

First we define the transformation that moves the keypoints at any given frame to the resident-centered coordinate frame. Let *t* index video frames and *k* = 1, …, *K* (*K* = 7) index keypoints. Let 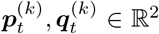 denote the resident and intruder keypoint coordinates in the arena frame, with centroids 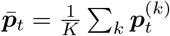 and 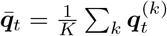. Finally, let

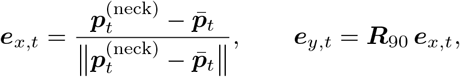

be the forward and leftward unit vectors. With these definitions, the heading angle *ϕ*_*t*_ = arg(***e***_*x,t*_) can be used to parameterize the arena-to-ego rotation ***Q***_*t*_ = ***R***(− *ϕ*_*t*_), and the egocentric transform can be defined as

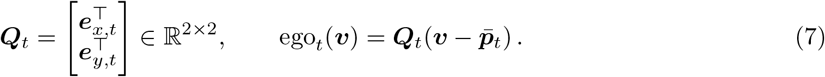

#### SELF: resident self-action features

Given the egocentric projection, the mapping of the resident’s position (or the single mouse for the single-mouse dataset) is defined as

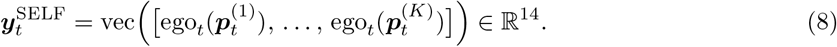

Note that with centroid centering as in Eq. (7), SELF satisfies 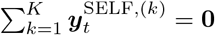.

#### DIST: dyadic distance features

Let 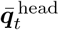 and 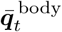 be the intruder head-keypoint centroid (nose, ears, neck) and body-keypoint centroid (neck, hips, tail base), and let ∥ · ∥ denote the Euclidean norm:

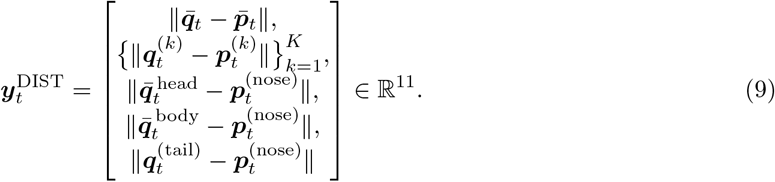

We further define *separation* as the average of the eleven distances 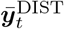.

#### DIRC: intruder direction features

The two components of the intruder direction features 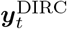 are the forward and leftward projections of the intruder centroid in the resident’s heading-aligned frame. Specifically these directions encode the intruder’s bearing relative to the resident’s heading:

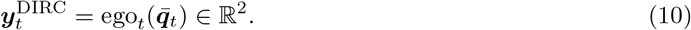

The magnitude of 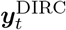 equals the centroid distance (the first entry of DIST), so that the information unique to DIRC is the angular direction ∠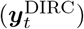, i.e., which way the resident would need to turn to face the intruder. To form the full feature set we concatenate these three sets of features in the concatenated feature vector ***y***_*t*_ ∈ ℝ^27^,

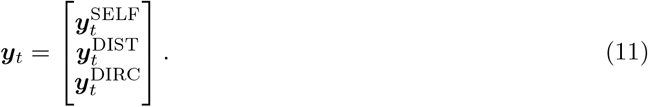

#### Post-encoding smoothing of ego-pose features

After egocentric transformation we apply a second Gaussian smoothing pass (*σ* = 0.5 frames) to the 14 SELF dimensions only. This smoothing mitigates effect where the ego-rotation, which can amplify small heading errors into pose jitter proportional to limb length. DIST and DIRC are not impacted by this effect and so do not require additional smoothing.

#### Egocentric orientation convention

For the egocentric directions we use the feature frame orientation as measured from above: +*x* denotes “forward” and +*y* = ***R***_90_***e***_*x,t*_ is “left”. We note that the source videos use image coordinates whose *y*-axis points *downward*. A turn or partner bearing toward +*y*, which we call *leftward*, can appear *rightward* in the raw video. For the single-mouse data, we use the same general construction, differing only in the heading axis and centering (§4.1).

### 4.3 Decomposed linear dynamical system

We fit dLDS, as described in §2.1, using the Julia dLDS package developed by [Koukuntla, 2025] that implements the standard dLDS update equations with the FISTA algorithm. Here we outline the model fitting details specific to our use of dLDS, including data-normalization, dictionary-fitting protocol, and the regularization terms used.

#### Data normalization

The raw features ***y***_*t*_ can differ in absolute scale, which can make model fitting hyper-focused on representing the features with the largest values. We thus standardized the features by normalizing to the temporal standard deviation and session-specific factors. Specifically, we define the normalized features ***x***_*t*_ = ***D***^*−*1^***y***_*t*_, where ***D*** is a diagonal matrix whose diagonal elements ***d***_*i*_ are

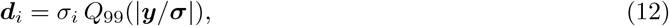

i.e., the product of the standard deviation of feature *i, σ*_*i*_, and the 99^*th*^ percentile of the normalized feature magnitudes, *Q*_99_(|***y****/****σ***|). To account for any inter-session variability, we computed and applied ***D*** on a per-session basis.

#### Dictionary fitting

We fit the dLDS dynamics operators via leave-one-session-out (LOSO) cross validation. To ensure the consistency of the operator signs (as there is an inherent sign-ambiguity in the model) we align the operator signs of each fold to the first fold. We use the first 30 folds to compute a consensus dictionary of operators 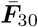, which we use as the dictionary set. With the final learned dictionary, we recompute all the signed coefficient trajectories for each sessions to obtain the operator activation time-courses. For the single-mouse data we fit just one dictionary jointly across all five sessions.

#### Regularization

To prevent the operators from capturing redundant dynamics, we introduce regularization ℛ into Eq. (4). Specifically, we regularize the pairwise correlations between the operators and the smoothness of the activations as

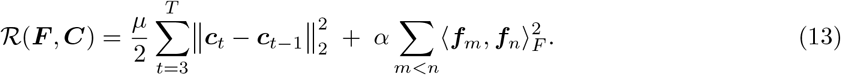

These regularization terms modify the dynamics operator update step as described in [Mudrik et al., 2024].

#### Hyperparameters

The hyperparameters we used for both datasets are provided in full in Table 1. We tested these parameters via a sensitivity analyses in the single-mouse dataset, varying the regularization *α*, sparsity parameter *λ*, and temporal smoothness parameter *µ* (Supp. Tab. S1b).

**Table 1:**
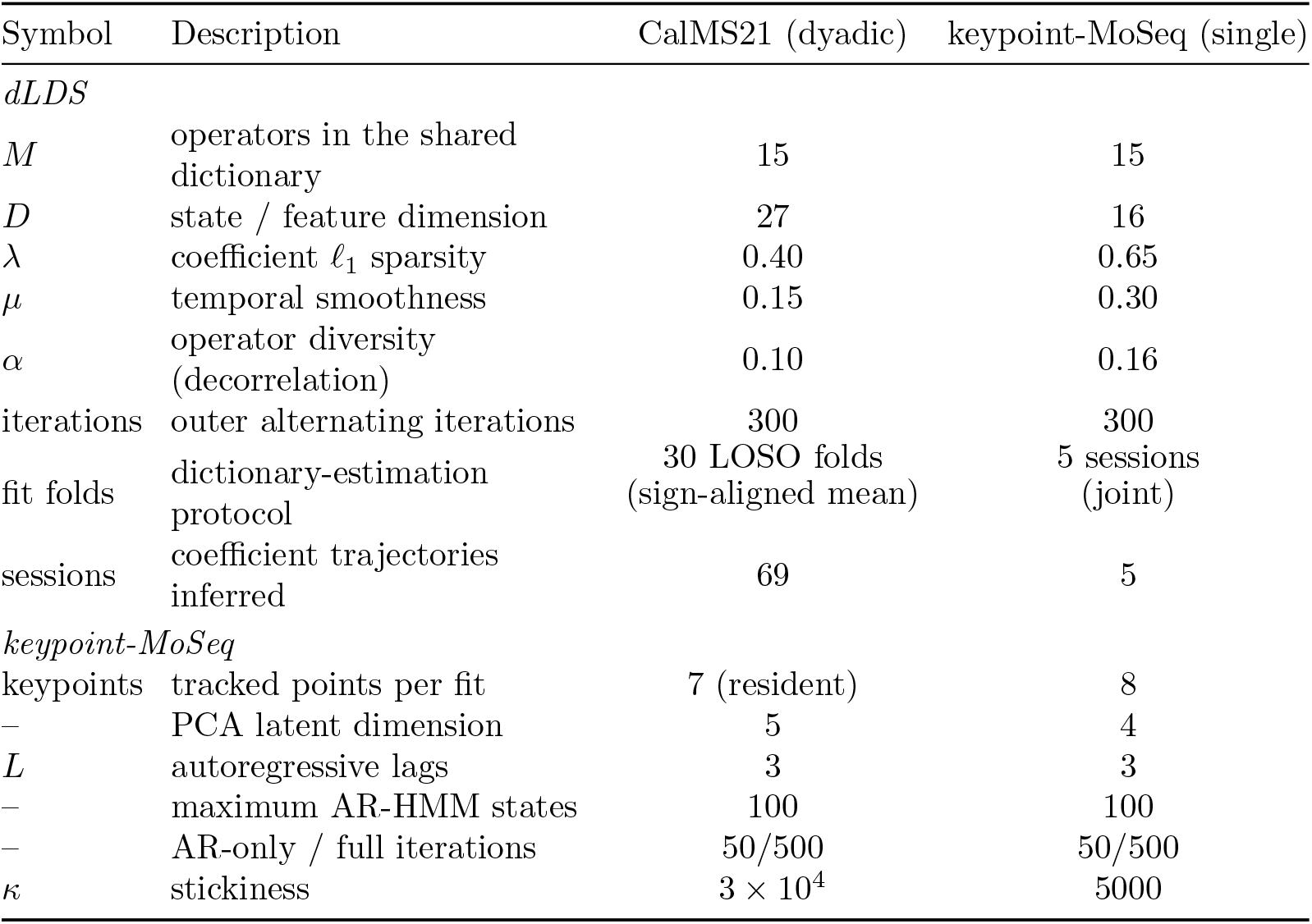
Model fitting settings for the two datasets.

| Symbol | Description | CalMS21 (dyadic) | keypoint-MoSeq (single) |
|---|---|---|---|
| *dLDS* |  |  |  |
| *M* | operators in the shared dictionary | 15 | 15 |
| *D* | state / feature dimension | 27 | 16 |
| λ | coefficient ℓ<sub>1</sub> sparsity | 0.40 | 0.65 |
| μ | temporal smoothness | 0.15 | 0.30 |
| α | operator diversity (decorrelation) | 0.10 | 0.16 |
| iterations | outer alternating iterations | 300 | 300 |
| fit folds | dictionary-estimation protocol | 30 LOSO folds<br>(sign-aligned mean) | 5 sessions<br>(joint) |
| sessions | coefficient trajectories inferred | 69 | 5 |
| *keypoint-MoSeq* |  |  |  |
| keypoints | tracked points per fit | 7 (resident) | 8 |
| – | PCA latent dimension | 5 | 4 |
| *L* | autoregressive lags | 3 | 3 |
| – | maximum AR-HMM states | 100 | 100 |
| – | AR-only / full iterations | 50/500 | 50/500 |
| κ | stickiness | 3 × 10<sup>4</sup> | 5000 |

### 4.4 Keypoint-MoSeq baseline

We fit keypoint-MoSeq [Weinreb et al., 2024] to both datasets with the released keypoint-moseq package. The parameter settings used for each dataset are provided in Table 1. We varied the value of *κ* in Supp. Tab. S1a to ensure an optimal fit to the data. We take as the main syllables those whose pooled usage is ≥ 0.5% (Supp. Tab. S1a).

### 4.5 Block-wise interpretation of operators

To interpret the learned operators, we first recognize that at each time-point *t*, the operator ***f***_*m*_ contributes to the next state ***x***_*t*_ the term *c*_*m,t*_ ***f***_*m*_ ***x***_*t−*1_. With the index sets defined in Eq. (11), we let ***f***_*m*_[*R, S*], with *R, S* ∈ {SELF, DIST, DIRC}, denote the submatrix that maps source block *S* at *t* − 1 to target block *R* at *t*. We can interpret each block by analyzing how it reads in a subset of the input sequences and how it projects those inputs onto the next time-point. For square blocks ***f***_*m*_[*R, R*] along the diagonal, we can perform this interpretation through the sub-matrix’s eigendecomposition. Specifically, the leading eigenvector represents the direction along which the given block projects the state and *λ*_1_ the corresponding per-step gain. For the rectang ular blocks ***f***_*m*_[*R, S*] with *R* ≠ *S*, we instead analyze the blocks using the singular value decomposition ∑_*i*_ *σ*_*i*_***u***_*i*_***v***_*i*_^*⊤*^. In this case the block projects the block of the state by projecting onto the right singular vector ***v***_*i*_, and then subsequently projects the result, scaled by the singular value, out into the next state through the right singular vector ***u***_*i*_. We found that for these interpretations, it was sufficient to consider only the top eigen- or singular-mode, as all blocks were well approximately rank-one.

#### Block profiles

Before interpreting any single block, we first identify for each operator which blocks are the primary drivers for that operator. We perform this identification by computing the mean absolute entry over each block, normalized over the nine blocks,

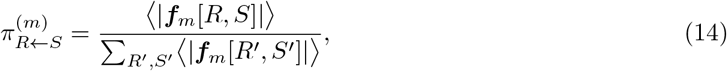

where ⟨| · |⟩ is the mean absolute entry of a block. The within-block averaging keeps larger blocks (e.g., SELF←SELF) from outweighing smaller ones (e.g., DIRC←DIRC).

#### Operator selection

For each retained operator, we next test whether its values are significantly concentrated within the blocks with highest mean-absolute values (boxes in Fig. 4d). The test compares the relative fraction of its mean-absolute value inside these blocks before and after randomly permuting the operator’s entries (shuffle test, §4.10). This shuffle enables us to report its *p*-value for significance of operator concentration within a given block.

#### Action and partner-geometry prediction

To understand if our operator interpretation was correct, we compared the action predicted from each operator with the action measured on the video frames in which the corresponding coefficient was non-zero (i.e., its “active” frames). We define the measured action as the mean deviation of the resident’s SELF features from their overall mean, 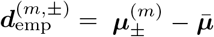, where 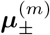 averages the SELF features at *t* over the active frames *t*, and 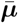 averages them over all frames. For the action predicted from the operator, we use the configurations provided from the operator restricted to the SELF block,

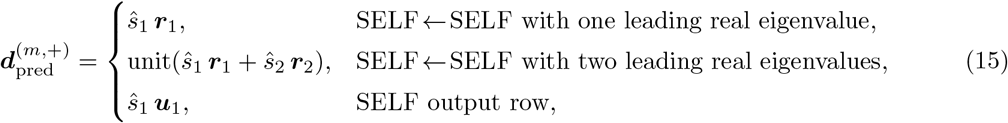

where ***r***_*k*_ is the real part of the *k*-th eigenvector of ***f***_*m*_[SELF, SELF] and ***u***_1_ is the leading left singular vector of the SELF output row ***f***_*m*_[SELF, :]. For the empirical (frame-driven) deviation, we compute the positive 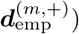 via its the projection with 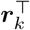, i.e., 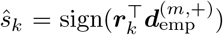, and set the negative version to be 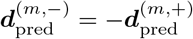. We define the agreement between predicted and empirical via

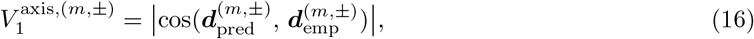

which we test for significance using a circular-shift null (§4.10).

### 4.6 Single-mouse operator–syllable correspondence

In the single-mouse model, the full state only consists of the 16-dimensional SELF block of the single animal (Tab. 1). We relate these operators, to keypoint-MoSeq syllables in two ways. First we compare the enrichment of syllables in each operator phase (§4.10). Second, we compare the subspace spanned by the operators’ predicted actions (§2.2).

#### Arena heading and angular velocity

The angular velocity of an animal is

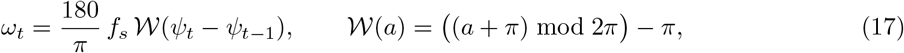

in degrees per second, with *f*_*s*_ = 30 Hz. Note that here *ω*_*t*_ *>* 0 indicates a left turn. The heading *ψ*_*t*_ is the angle from spine4 to the nose. We compute the angular velocity of a syllable as the mean of *ω*_*t*_ over all the frames where that syllable is active, pooled over all sessions.

#### Shared operator subspaces

To analyze the shared subspace between dLDS operators and syllables, we define a shared subspace using predicted actions of the three dLDS operators. We convert each ***d***_pred_ (Eq. (15)) from the normalized state to pose units by multiplying each feature by its standard deviation over all frames. We then orthonormalize the three directions with a QR decomposition to obtain ***U***_op_∈ ℝ ^16*×*3^. For the syllables *S* with usage ≥0.5% (§4.4), let ***µ***_*s*_ be the mean pose of syllable *s* and 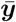 the mean pose over all frames. We compute the share of the syllables’ mean deviations that lies in the operator subspace as

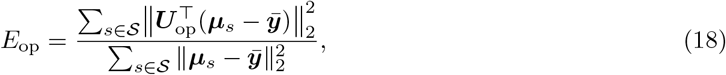

where the summation sums over all syllables.

#### Alignment with the syllable representation space

To embed syllables into the representation space we look at how ***U***_op_ aligns with the principle direction of the syllables. Let **Δ** ∈ ℝ ^|*S*|*×*16^ be the matrix formed by vertically concatenating the deviations 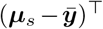, and let ***V*** = [***v***_1_ ***v***_2_ ***v***_3_] represent the top three right singular vectors of **Δ**. We can then compute the principal angles between these two subspaces via the inverse cosines of the singular values of 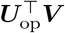. For reference, we compared the resulting principle angles to those computed for 2,000 random three-dimensional subspaces. To further assess how well the found three dimensional subspace embedded the data, we computed how much of each syllable’s action lies outside span(***V***) via

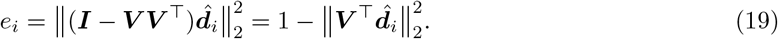

### 4.7 Partner geometry and kinematics

#### Trunk axes, relative heading and bearing

The trunk axis of each animal runs from its centroid to its neck. For the resident this is the forward axis ***e***_*x,t*_ with angle *ϕ*_*t*_ (§4.2). 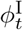 is the angle of the intruder’s trunk axis. We compute the relative heading as 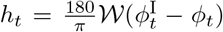, with W as in Eq. (17). In this definition, *h*_*t*_ *>* 0 indicates that the intruder’s trunk axis points to the resident’s left (and similarly for *h*_*t*_ *<* 0 indicating right). We group *h*_*t*_ into four 90^*°*^ categories, “same” (−45^*°*^ ≤ *h*_*t*_ *<* 45^*°*^), “left” (45^*°*^ ≤ *h*_*t*_ *<* 135^*°*^), “right” (−135^*°*^ ≤ *h*_*t*_ *<* −45^*°*^) and “opposite” (135^*°*^ ≤ *h*_*t*_ *<* 180 and −180 *< h*_*t*_ *<* −135^*°*^). For a set of angles {*θ*_*t*_}, we compute the circular mean as arg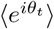 and the mean resultant length is 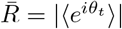.

#### Kinematics

To compute the velocities of each mouse we project the temporal difference between consecutive keypoint states through the operator/syllable. In this way, the velocities of the two centroids are computed as 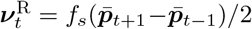 and 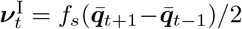 in pixels per second, with *f*_*s*_ = 30Hz. Defining the unit vector ***ê***_*t*_ from 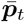 to 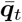, the resident’s chase speed is 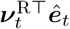 and the intruder’s flee speed is 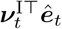. The flee speed minus the chase speed thus represents the rate of change of the inter-animal distance 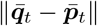.

### 4.8 Temporal dynamics

#### Behavior bouts

To relate operator activity to annotated behavior, we align the coefficients on behavior bouts. We define a bout as a consecutive run of one behavior label of at least 0.5 s. We call the first frame of each bout as the “bout onset” and the first frame after the bout as the “bout offset”.

#### Pulses of *c*_9_

Because *c*_9_ rises and falls repeatedly within a mount bout, we divide the time series into pulses at the troughs between the coefficient peaks. Within each session, we find the positive peaks of *c*_9_ that have a prominence of at least 0.02 and lie at least 5 frames apart. Between two consecutive peaks, we take the frame where |*c*_9_| is smallest as the trough. When the minimum spans several frames, we take the middle frame within that span as the trough. After identifying pulses, we measure the length of each pulse (trough-to-trough) and only keep pulses with duration 0.3 − 1.2 s, and filter out any pulse with peak value *<* 0.05 or have frames within the pulse that are not “near” frames. To average pulses of different lengths, we map the rise onto [0, 0.5] and the fall onto [0.5, 1], and interpolate each pulse onto 41 evenly spaced points on this axis.

#### Attack events

During the attack bouts, *c*_15_ is active in brief, isolated bursts rather than the more sustained, oscillatory nature of *c*_9_’s mount dynamics. We thus instead treat each burst as one event. We specify an event as a sequence of contiguous frames where *c*_15,*t*_ *>* 0 and whose frames are all labeled “attack”. We align the events to the peak (largest value of *c*_15_ in that event). Similar to the mount condition for *c*_9_, we keep an event only when this peak value is ≥ 0.05.

### 4.9 Behavior Decoding

#### Classifier details

We train behavioral decoders (classifiers) in a one-against-all setting where we seek to decode a single social behavior against all others. To classify each individual frame at time *t*, we use either the signed coefficients (for dLDS) or the one-hot keypoint-MoSeq syllable. Given these features, we train a logistic regression classifier, with class weights inversely proportional to the class frequencies. For each behavior, we use the sessions with enough frames, given the relative rarity of each behavior. For attack, we require at least 0.5% of frames to have the “attack” label, 1% for mount, and 2% for investigation. All classifiers are evaluated via leave-one-session-out cross-validation. To compute classifier performance on a given feature set, we compute the mean of the held-out session area-under-the-curve (AUCs), normalized by 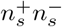, the product of the number of frames in session *s* labeled with that behavior and the number of frames with other different labels. To compute an error distribution for our classifiers, we repeat the fit over 200 resamples of the sessions with replacement, and compute the standard deviation over these resamples.

#### Greedy-minimum ladder

We aim to show how each operator contributes to the behavior decoding in order to build intuition for the behavior-related information in each operator. To accomplish this we start from a decoder that uses the single operator with the lowest AUC of all operators and add, at each step, the next operator that gives the minimal improvement to AUC. We add operators in this way until all eight operators are included.

For keypoint-MoSeq, which has many more syllables than dLDS has operators, we instead split the analysis into two decoders: one trained only on the most relevant syllables, as measured through their enrichment to that behavior, and another using all 28 syllables. For “attack”, we compared the “attack”-enriched decoder and 28-syllable decoder with the dLDS decoder using all eight operators by a paired bootstrap over sessions, and report its 95% interval, to see the results’ consistency, since “attack” is only active on 17 sessions. Note that all comparisons for each behavior share the same resamples.

### 4.10 Statistical tests

#### Circular-shift null

Operator coefficients and behavior labels change slowly, so neighboring frames are not independent samples. We therefore test each statistic computed from the time series against circular shifts of that series in order to not break the temporal correlations. Specifically, we produce *N* = 1,000 replicates, where each replicate consists of a shift of the series by a lag drawn uniformly from 1 to *T*_*s*_ − 1, where *T*_*s*_ is the session length. The state, labels, and all other time-series are not shifted (thereby providing the necessary shuffle), and we recompute the statistic of interest on these relatively shifted series. In this shuffle test, the *p*-value is *p* = (*n* + 1)*/*(*N* + 1), where *n* is the number of replicates whose statistic is at least as large as the one computed on the original data. Three types of statistics use this shuffle test: the enrichment of behaviors or syllables on the operator phases; the resident’s action prediction *V*_1_ (Eq. (16)); and the intruder’s position prediction (Supp Fig. S3a).

#### Shuffle test

To identify if the concentration of an operator on a specific set of blocks (Operator selection, §4.5) is significant, we perform a shuffle test over the entries of the matrix. to do so we generate *N* = 10,000 matrices with the entries of the matrix ***f***_*m*_ are randomly permuted and recompute the relative concentration of ***f***_*m*_ in the identified blocks. Under this shuffle test we can similarly compute the *p*-value as above: *p* = (*n* + 1)*/*(*N* + 1).

## Data availability

The single-mouse data are available from Zenodo at https://doi.org/10.5281/zenodo.15171335, and the CalMS21 dyadic data are available from CaltechDATA at https://doi.org/10.22002/D1.1991. The fit dLDS operators and coefficient arrays used in this study are available at https://drive.google.com/file/d/1wBcfj0d4gs-eSbJVkumU--NDH6VPbbH8/view?usp=drive_link.

## Code availability

Code for data preprocessing, dLDS fitting, statistical analysis, and generation of the data-driven figure panels is available at https://github.com/YueWan1/social-dLDS.

## Acknowledgments

Y.W. was supported by a Johns Hopkins Discovery Award. A.S.C. was supported by NSF CAREER Award

## Appendix A Supplementary Notes

### Supplementary tables and figures

**Table S1:**
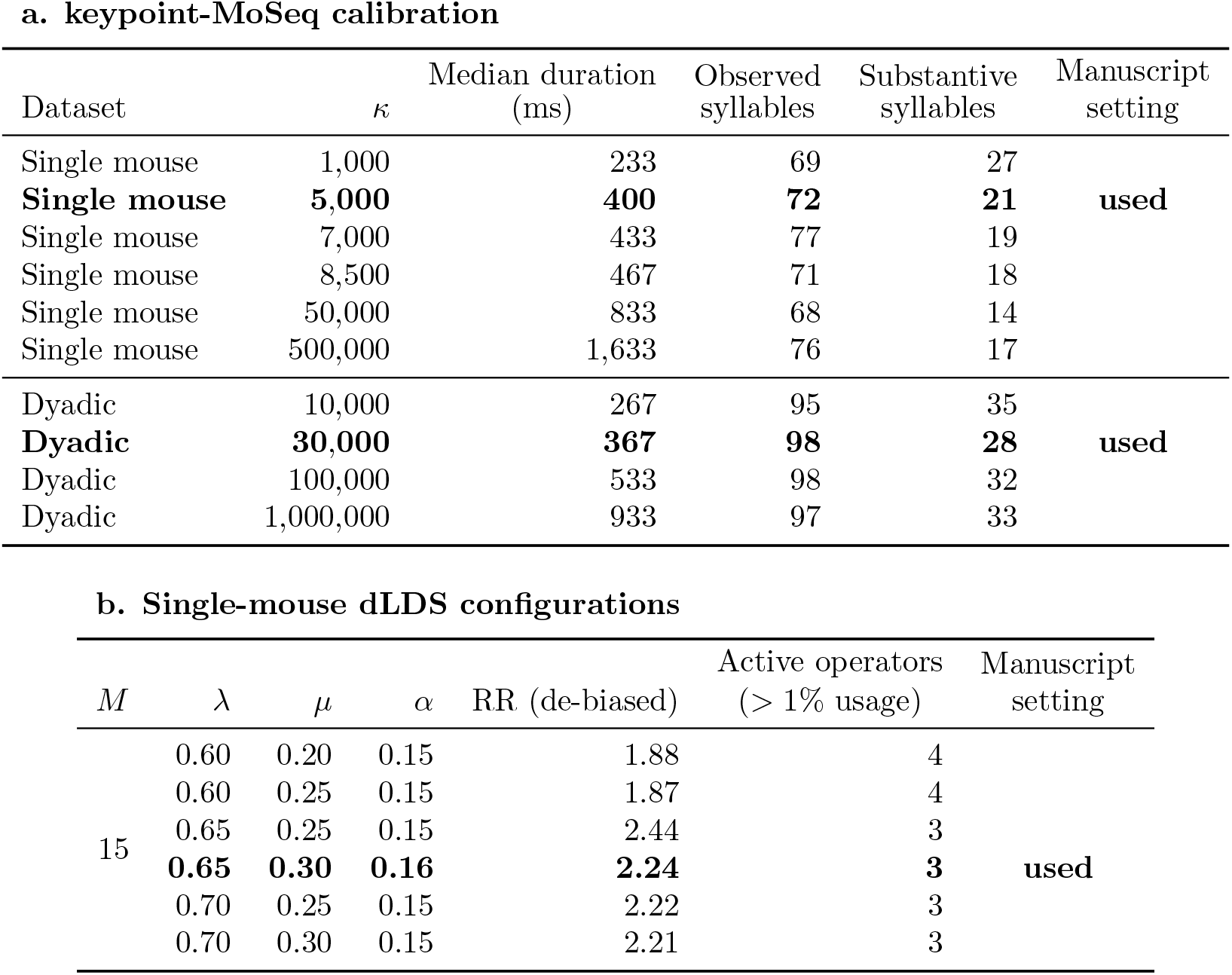
Sensitivity of model granularity across the tested parameter settings. **a**, keypoint-MoSeq temporal calibration; substantive syllables have pooled usage ≥ 0.5%. **b**, Single-mouse dLDS configurations (*M* = 15, 300 iterations).

| **a. keypoint-MoSeq calibration** |  |  |  |  |  |  |
| Dataset | κ | Median duration (ms) | Observed syllables | Substantive syllables | Manuscript setting |  |
|---|---|---|---|---|---|---|
| Single mouse | 1,000 | 233 | 69 | 27 |  |  |
| **Single mouse** | **5,000** | **400** | **72** | **21** | **used** |  |
| Single mouse | 7,000 | 433 | 77 | 19 |  |  |
| Single mouse | 8,500 | 467 | 71 | 18 |  |  |
| Single mouse | 50,000 | 833 | 68 | 14 |  |  |
| Single mouse | 500,000 | 1,633 | 76 | 17 |  |  |
| Dyadic | 10,000 | 267 | 95 | 35 |  |  |
| **Dyadic** | **30,000** | **367** | **98** | **28** | **used** |  |
| Dyadic | 100,000 | 533 | 98 | 32 |  |  |
| Dyadic | 1,000,000 | 933 | 97 | 33 |  |  |
| **b. Single-mouse dLDS configurations** |  |  |  |  |  |  |
| *M* | λ | μ | α | RR (de-biased) | Active operators (> 1% usage) | Manuscript setting |
| 15 | 0.60 | 0.20 | 0.15 | 1.88 | 4 |  |
|  | 0.60 | 0.25 | 0.15 | 1.87 | 4 |  |
|  | 0.65 | 0.25 | 0.15 | 2.44 | 3 |  |
|  | **0.65** | **0.30** | **0.16** | **2.24** | **3** | **used** |
|  | 0.70 | 0.25 | 0.15 | 2.22 | 3 |  |
|  | 0.70 | 0.30 | 0.15 | 2.21 | 3 |  |

**Table S2:**
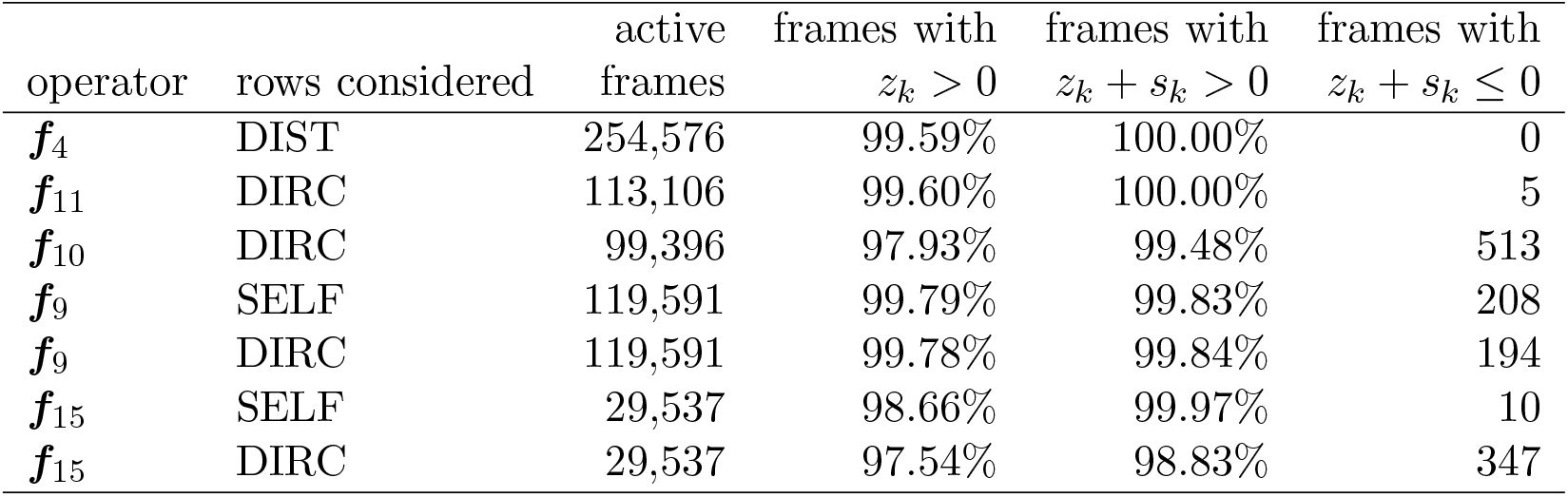
Sign of the input scalars. Sign of the projection of the state onto the right singular/eigen vector on the frames where each operator is active.

|  |  | active | frames with | frames with | frames with |
|---|---|---|---|---|---|
| operator | rows considered | frames | *z*<sub>*k*</sub> > 0 | *z*<sub>*k*</sub> + *s*<sub>*k*</sub> > 0 | *z*<sub>*k*</sub> + *s*<sub>*k*</sub> ≤ 0 |
| ***f***<sub>4</sub> | DIST | 254,576 | 99.59% | 100.00% | 0 |
| ***f***<sub>11</sub> | DIRC | 113,106 | 99.60% | 100.00% | 5 |
| ***f***<sub>10</sub> | DIRC | 99,396 | 97.93% | 99.48% | 513 |
| ***f***<sub>9</sub> | SELF | 119,591 | 99.79% | 99.83% | 208 |
| ***f***<sub>9</sub> | DIRC | 119,591 | 99.78% | 99.84% | 194 |
| ***f***<sub>15</sub> | SELF | 29,537 | 98.66% | 99.97% | 10 |
| ***f***<sub>15</sub> | DIRC | 29,537 | 97.54% | 98.83% | 347 |

**Table S3:** Computational cost of single-mouse model. Timing and peak memory running on dLDS and keypoint-MoSeq on one 24-core machine no other processes competing for resources.

|  | keypoint-MoSeq | dLDS | keypoint-MoSeq / dLDS |
|---|---|---|---|
| start-up: interpreter, packages, workers (s) | 2.27 | 8.86 |  |
| *Fitting* |  |  |  |
| load/normalize model input (s) | - | 0.90 |  |
| model initialization (s) | 5.50 | - |  |
| stage 1: AR-HMM, 51 sweeps (s) | 107.4 | - |  |
| stage 2: 501 sweeps / 300 iterations (s) | **3637.9** | **77.60** | 46.9× |
| inference over all 375 809 frames (s) | 0.01 | 2.33 | 0.004× |
| **total, model input to fitted model** (s) | **3753.2** | **91.3** | **41.1**× |
| in minutes | 62.6 | 1.52 |  |
| peak memory (GB) | 5.71 | 8.76 | 0.65× |
| cores used during fitting (of 24) | 6.64 | 7.81 |  |

**Table S4:** Computational cost as a function of CPU cores allocated. Same sessions and machine as Table S3. Each dLDS run reproduced the same result to machine precision.

| Cores | keypoint-MoSeq |  |  |  |  | dLDS |  |  |  |
|---|---|---|---|---|---|---|---|---|---|
| allocated | AR | full | fitting | speedup | used | per iter | fitting | speedup | used |
|  | (s/sweep) |  | (s) |  |  | (s) | (s) |  |  |
| 1 | 3.357 | 11.722 | 6055 | 1.00 | 1.00 | 1.285 | 392.2 | 1.00 | 1.00 |
| 4 | 2.380 | 7.921 | 4093 | 1.48 | 2.29 | 0.556 | 170.0 | 2.31 | 2.64 |
| 12 | 2.142 | 7.347 | 3792 | 1.59 | 4.19 | 0.258 | 79.7 | 4.98 | 7.83 |
| 24 | 2.146 | 7.268 | 3753 | 1.61 | 6.67 | 0.228 | 70.5 | 5.65 | 14.06 |

**Table S5:** Validation of resident action and intruder bearing prediction. Action prediction and intruder direction are tested using the circular-shift null (Methods 4.10).

| Phase | ***d***<sub>pred</sub> | Active frames | \|cos\| | *p* | Bearing error (Δ°) |
|---|---|---|---|---|---|
| ***f***<sub>2</sub><sup>+</sup> | ***v***<sub>1</sub> + ***v***<sub>2</sub> | 129,227 | 0.970 | 0.001 |  |
| ***f***<sub>2</sub><sup>−</sup> | ***v***<sub>1</sub>, ***v***<sub>2</sub> | 8,869 | 0.934 | 0.001 |  |
| ***f***<sub>3</sub><sup>+</sup> | ***v***<sub>1</sub> | 456,604 | 0.982 | 0.001 |  |
| ***f***<sub>6</sub><sup>+</sup> | ***v***<sub>1</sub> + ***v***<sub>2</sub> | 146,633 | 0.965 | 0.001 |  |
| ***f***<sub>6</sub><sup>−</sup> | ***v***<sub>1</sub>, ***v***<sub>2</sub> | 21,973 | 0.847 | 0.009 |  |
| ***f***<sub>9</sub><sup>+</sup> | +***u***<sub>9</sub> | 58,935 | 0.923 | 0.001 | 5.4° |
| ***f***<sub>9</sub><sup>−</sup> | −***u***<sub>9</sub> | 60,656 | 0.814 | 0.169 | 28.3° |
| ***f***<sub>15</sub><sup>+</sup> | +***u***<sub>15</sub>, ***u***′<sub>15</sub> | 15,286 | 0.733 | 0.001 | 3.9° |
| ***f***<sub>15</sub><sup>−</sup> | −***u***<sub>15</sub>, −***u***′<sub>15</sub> | 14,251 | 0.603 | 0.001 | 2.3° |

**Table S6:** One-step reconstruction errors for different keypoints. All errors reported in units of pixels. Aggregate errors reported with the Relative Reconstruction Error 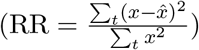, M is dictionary size, and “Used” indicates the number of prominent operators used by the model.

|  | *M* | Used | RR (de-biased) |
|---|---|---|---|
| dLDS, ℓ<sub>1</sub> = 0.30 | 10 | 6 | 1.59 |
| dLDS, ℓ<sub>1</sub> = 0.40 | 10 | 5 | 1.43 |
| dLDS, ℓ<sub>1</sub> = 0.55 | 10 | 5 | 1.65 |
| dLDS, ℓ<sub>1</sub> = 0.60 | 15 | 4 | 1.87 |
| **dLDS, ℓ<sub>1</sub> = 0.65** | **15** | **3** | **2.24** |
| dLDS, ℓ<sub>1</sub> = 0.70 | 15 | 3 | 2.21 |
| keypoint-MoSeq |  |  | 1.11 |

**Table S7:**
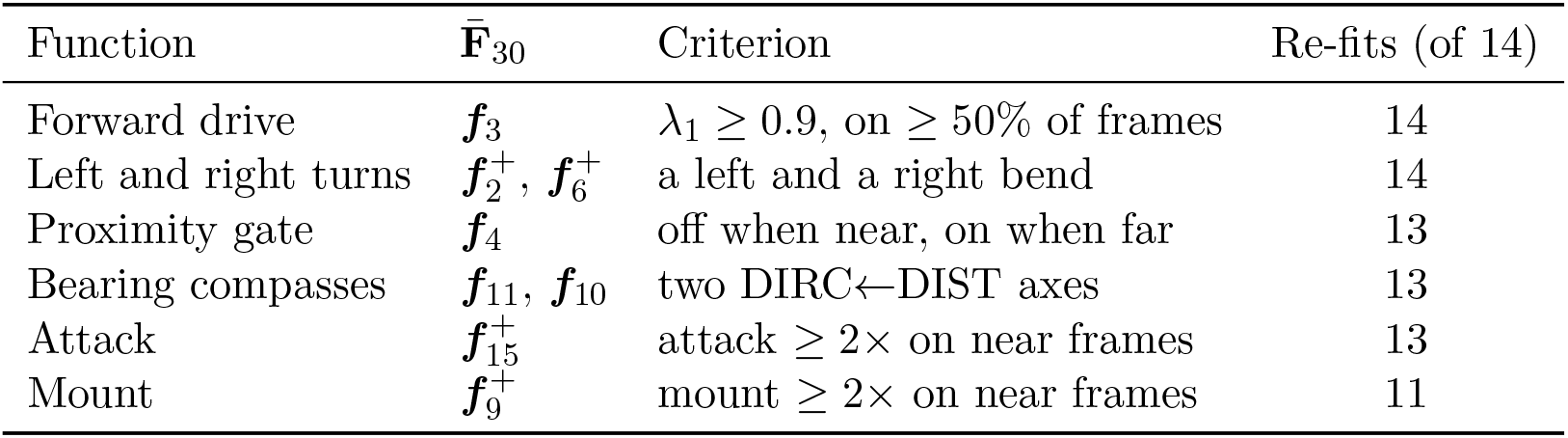
Recurrence of the operator functions of 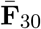 in 14 reinitializations. 10/14 completely replecated all operators including mount and attack.

| Function | **F̄**<sub>30</sub> | Criterion | Re-fits (of 14) |
|---|---|---|---|
| Forward drive | ***f***<sub>3</sub> | λ<sub>1</sub> ≥ 0.9, on ≥ 50% of frames | 14 |
| Left and right turns | ***f***<sub>2</sub><sup>+</sup>, ***f***<sub>6</sub><sup>+</sup> | a left and a right bend | 14 |
| Proximity gate | ***f***<sub>4</sub> | off when near, on when far | 13 |
| Bearing compasses | ***f***<sub>11</sub>, ***f***<sub>10</sub> | two DIRC←DIST axes | 13 |
| Attack | ***f***<sub>15</sub><sup>+</sup> | attack ≥ 2× on near frames | 13 |
| Mount | ***f***<sub>9</sub><sup>+</sup> | mount ≥ 2× on near frames | 11 |

**Figure S1:**
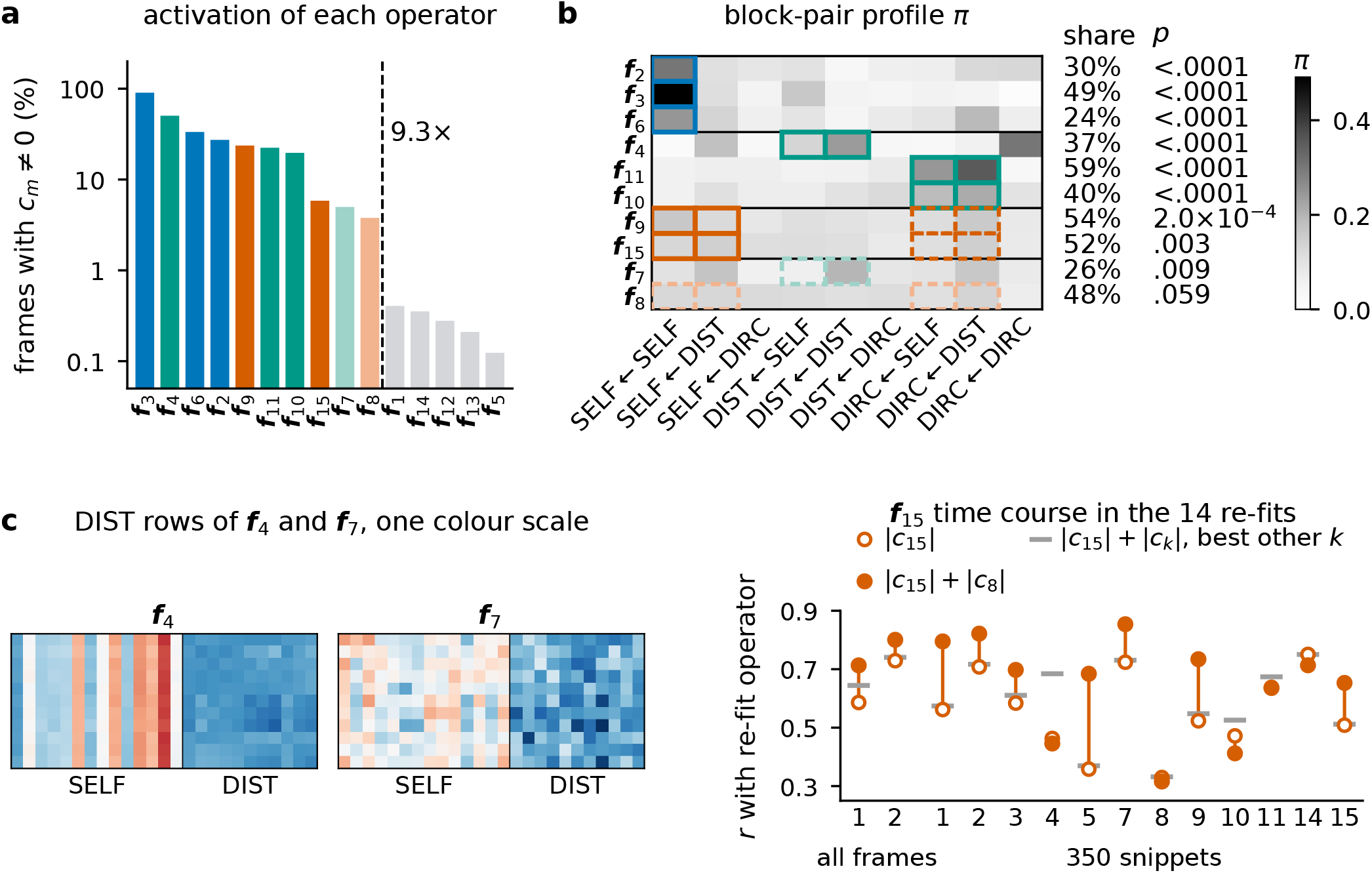
Operator selection for the dyadic dictionary. **a)** Share of frames for which each operator is active (*c*_*m,t*_ ≠ 0). The dashed line marks the largest gap, which represents a 9.3× drop in the ranking. **b)** Block-wise concentration profile (Eq. (14)) of the ten engaged operators. Solid boxes mark the blocks with the highest concentration that we use to analyze the operators. Right, percent of operator values inside the boxes. *p*-values are computed with respect to 10,000 permutations of the operator’s entries. **c)** Left, the DIST rows of ***f***_4_ and ***f***_7_. Right, consistency of attack operator. We refit the dictionary 14 times with different random seeds and compute the correlation between the absolute magnitude of the coefficients |*c*| of the operator that best matches ***f***_15_ and 1) |*c*_15_| (the open circles), |*c*_15_| + |*c*_8_| (filled circles), or |*c*_15_| plus the other coefficient that most increases the correlation (grey lines).

**Figure S2:**
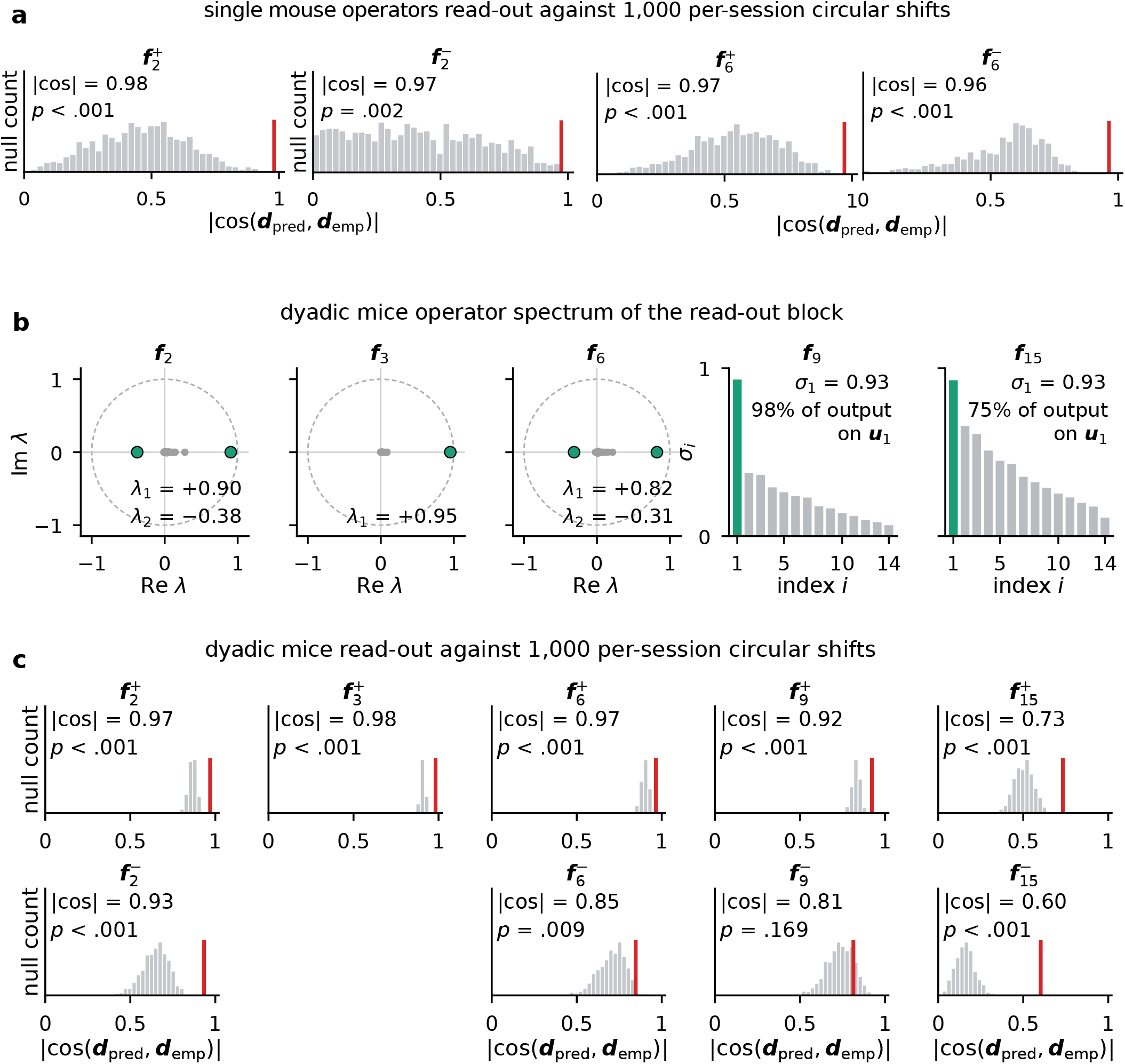
Within-block and cross-block interpretation of both datasets operators. **a)** Single mouse operators (±***f***_2_, ±***f***_6_),| cos(***d***_pred_, ***d***_emp_)| (red line) as compared to 1,000 per-session circular shifts of *c* (grey). All five phases exceed their nulls, we didn’t test ***f***_14_ because nearly all frames of ***f***_14_ are positive (47 frames exception). **b)** EVD of each SELF←SELF block (for ***f***_2_, ***f***_*w*_, and ***f***_6_; left three plots), and the SVD for the full SELF rows (for ***f***_9_ and ***f***_15_; right two plots). **c)** Dyadic operators | cos(***d***_pred_, ***d***_emp_)| (red line) as compared to 1,000 per-session circular shifts of *c* (grey). All nine phases exceed their nulls.

**Figure S3:**
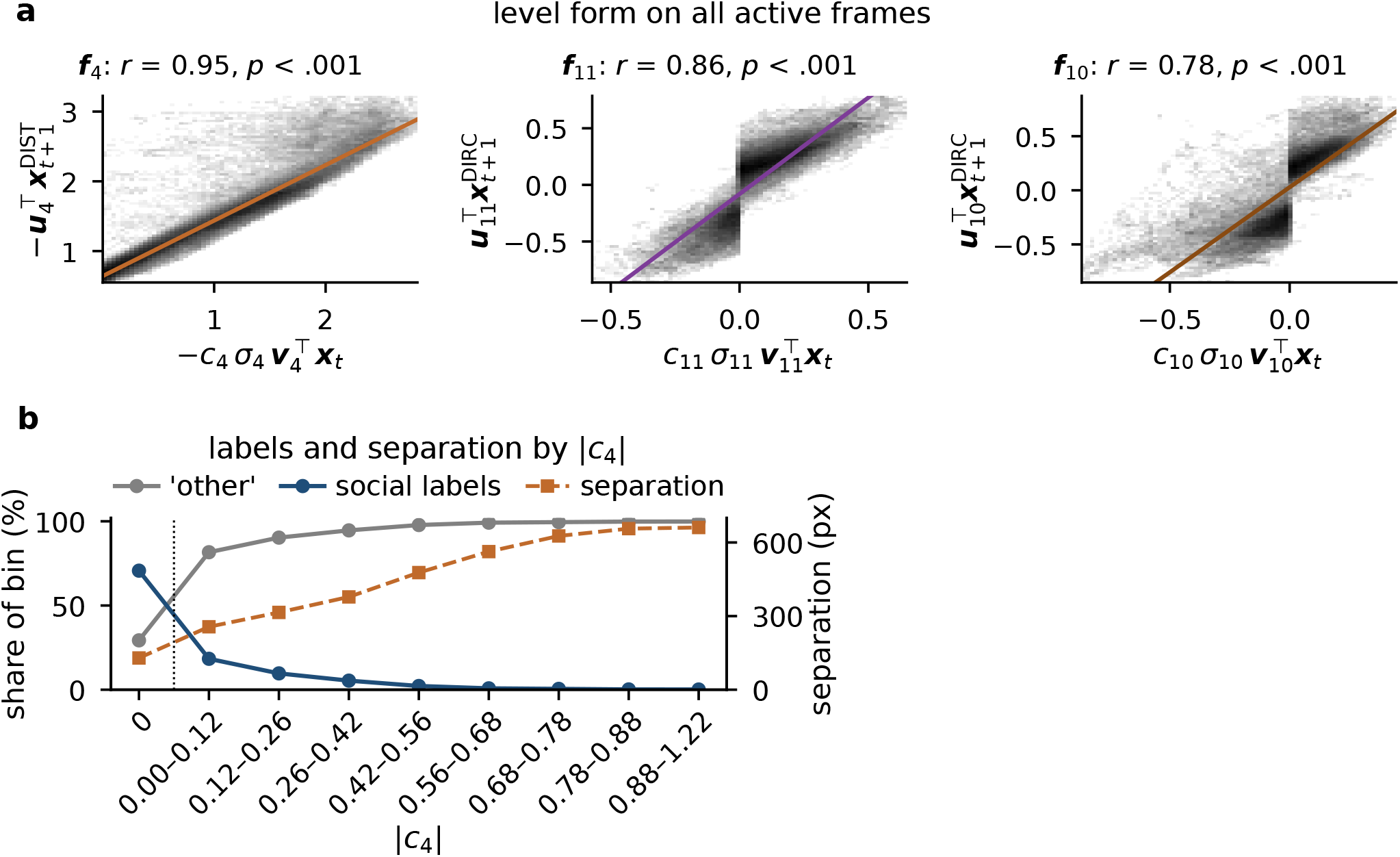
Additional validation of *f*_4_ and *f*_11_*/f*_10_ as tracking intruder separation and orientation. **a)** Comparison of the predicted position and actual position as projected into the top singular mode of the most relevant operator blocks. The x-axes show the predicted position (either DIST or DIRC) as computed by the projection of the state at time *t*, scaled by the singular value and operator coefficient *c*. The y-axes compute the magnitude of the projection of the left singular vector with the next time-step’s state. High correlations in the Pearson correlation *r* indicate that the operator is highly predictive of the state features determined by the block-wise analysis. We validate this analysis with 1,000 per-session circular shifts of *c*_*k*_, using the position-prediction test (see Methods. 4.10). **b)** Share of “other” and of “attack”, “investigation” and “mount” in bins of |*c*_4_| (*c*_4_ = 0, then eight bins holding equal numbers of frames), and the median separation of each bin (mean of the eleven distances; right axis).

**Figure S4:**
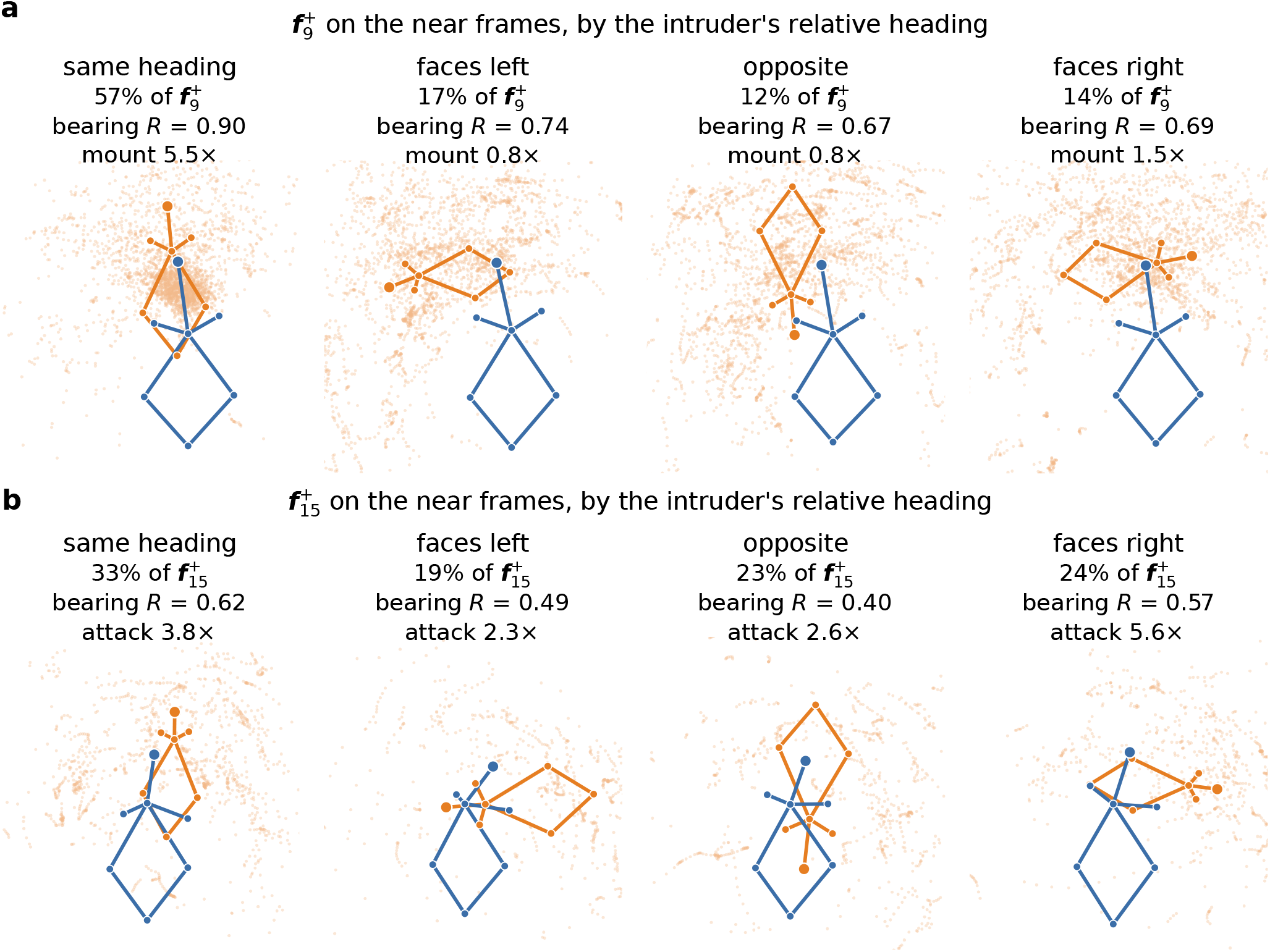
Aggregate orientations of the resident and intruder during *f*_9_ and *f*_15_ activations. **a)** poses during 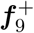 events, conditioned on the “near” frames and grouped by the intruder’s heading relative to the resident. The resident (blue) and intruder (orange) are depicted in their average raw-keypoint poses. Each panel shows that orientation’s share of the total 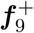 frames, the *R* of the intruder’s bearing, and its mount enrichment over all “near” frames. **b)** The same as a), but for 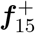 and “attack”.

